# The essential host genome for *Cryptosporidium* intracellular survival exposes metabolic dependencies that can be leveraged for treatment

**DOI:** 10.1101/2024.12.04.626561

**Authors:** N. Bishara Marzook, Ok-Ryul Song, Lotta Baumgärtel, Netanya Bernitz, Tapoka T. Mkandawire, Lucy C. Watson, Vanessa Nunes, Scott Warchal, James I. MacRae, Michael Howell, Adam Sateriale

**Affiliations:** The Cryptosporidiosis Laboratory, The Francis Crick Institute, UK; High-throughput Screening Platform, The Francis Crick Institute, UK; Metabolomics Science Technology Platform, The Francis Crick Institute, UK

**Author notes:** Correspondence: Adam Sateriale.

## Abstract

Mapping how pathogens interact with their host cells can reveal unexpected pathogen and host cell biology, paving the way for new treatments. *Cryptosporidium* is an intracellular parasite of intestinal epithelial cells, and a leading cause of diarrheal death and disease in infants worldwide. Despite this, very little is known about the cell biology of infection of this eukaryotic pathogen. Here, we designed and implemented a unique microscopy-based arrayed CRISPR-Cas9 screen to interrogate the effects of the loss of every protein-coding human gene on a *Cryptosporidium* infection. As the experimental readout is image-based, we extracted multiple phenotypic features of infection, including parasite growth, progression of the parasite to its sexual life stage, and recruitment of host actin to ‘pedestals’ beneath the parasite vacuole. Using this dataset, we discovered a tipping point in the host cholesterol biosynthesis pathway that controls *Cryptosporidium* infection. Parasite growth can either be inhibited or promoted by the intermediary metabolite squalene. A build-up of squalene in epithelial cells creates a reducing environment, with more reduced host glutathione available for uptake by the parasite. Because *Cryptosporidium* has lost the ability to synthesise glutathione, this uptake from the host cell is required for growth and progression through its life cycle. We demonstrate that this dependency can be leveraged for treatment with the abandoned drug lapaquistat, an inhibitor of host squalene synthase that has efficacy against *Cryptosporidium in vitro* and *in vivo*.

## INTRODUCTION

*Cryptosporidium* is an obligate intracellular parasite of the gut. Within the environment, the parasite exists in a dormant, but infectious, state known as an oocyst. Upon ingestion, this oocyst releases motile forms of the parasite that invade epithelial cells that line the intestine. Here, the parasite creates an apically-localised vacuole, with a host actin-derived “pedestal” directly underneath. Inside this vacuole, the parasite will asexually replicate to exponentially infect neighbouring epithelial cells. After multiple rounds of asexual replication, parasites enter the sexual stage of their life cycle, transforming into females (macrogamonts) or males (microgametes) that need to find each other to produce new infectious oocysts^1,2^. Unlike other related apicomplexan parasites, such as *Plasmodium* and *Toxoplasma*, *Cryptosporidium* completes both asexual and sexual stages of its life cycle in a singular host, making it a unique system to better understand how host cell biology influences the parasite life cycle. Despite this, very little is known about the cell biology of infection of *Cryptosporidium*: what are the host factors that allow (or that are manipulated to allow) the parasite to invade, replicate, and survive within intestinal epithelial cells?

A growing appreciation of *Cryptosporidium*’s significant contribution to the burden of diarrhoeal-related deaths and disease in infants from lower-middle income regions ^3–6^, the immunocompromised^7^, and commercial livestock^8^ has recently spurred more active research into this parasite, particularly in the search for new therapeutics^9^. This urgency is particularly pronounced as the only FDA-approved drug for cryptosporidiosis, nitazoxanide, has proved ineffective in vulnerable populations^10^. We reasoned that focusing on the parasite’s host dependencies would reveal unexplored insights into the cell biology of infection of this deadly parasite, while also uncovering potentially novel therapeutic avenues to improve disease outcomes. To that end, we devised a unique microscopy-based arrayed CRISPR screen to systematically examine how every human gene influences a *Cryptosporidium* infection. In previously described host cell-directed CRISPR screens, Cas9-expressing cells are first transduced in bulk with a pool of guide RNAs. This mixed population of host cells is then infected by the pathogen of interest and surviving cells are sequenced to provide relative abundances of modified host cells before and after the assault^11–14^. While powerful and broadly applicable, this approach only reveals one dimension of the infection, specifically, which genes impact host cell survival. In the arrayed system presented here, each host gene is ablated individually, and the subsequent pathogen infection is monitored by high-content imaging and automated image analysis pipelines. This results in a much richer dataset where the impact of host gene loss can be examined across many parameters of infection.

In our analysis, the parameters of *Cryptosporidium* infection that we primarily focused on were parasite growth, parasite sexual development, host cell viability, and host actin recruitment to parasite vacuoles. We show that while the loss of certain host genes and pathways increased parasite survival, others had an inhibitory effect. Curiously, perturbation of one host metabolic pathway – cholesterol biosynthesis – appeared to have opposing effects on parasite growth and development. On closer analysis, we discovered the ability of squalene, an intermediate metabolite in the cholesterol biosynthesis pathway, to control parasite survival and development. Although *Cryptosporidium* does not appear to import and utilise host squalene, we found the accumulation of this metabolite strongly influences reactive oxygen species (ROS) levels and glutathione stores within intestinal epithelial cells. Further, the *Cryptosporidium* parasite appears to have lost the ability to synthesise its own glutathione, becoming reliant on the stores within the infected host cell. This has created a host-targeted Achilles heel that we demonstrate can be exploited for treatment.

## RESULTS

### A robust arrayed CRISPR-Cas9 screen for deconvoluting host-pathogen interactions

*In vitro, Cryptosporidium* completes multiple asexual replication cycles and then commits to sexual development over a 48-hour period. This developmental trajectory from asexual to sexual stages is thought to be necessary for the completion of this parasite’s life cycle^2,15^. To map *Cryptosporidium*’s interactions with, and dependencies on, its host cell over this period, we devised a microscopy-based arrayed full-genome CRISPR screen to enable us to analyse several parameters of infection during this complex life cycle. For a detailed description of how we designed and conducted the screen, please see the Methods section. Briefly, intestine epithelial cells (HCT-8) expressing Cas9 were seeded into clear-bottomed 384-well plates, where each well contained four guide RNAs targeting a single host gene. This was done for more than 18,000 human genes, covering virtually all human protein coding genes in triplicate. Cells were then infected with *C. parvum* parasites and fixed 49 hours post-infection (Fig 1A). Plates were prepared for immunofluorescence microscopy with fluorophores marking parasite vacuoles, female parasites, host actin, and host nuclei (Fig S1E). Each plate was subjected to high-content confocal microscopy to extract several image features from each channel, for each well (see Table S1 for a full list). Therefore, for every host gene knocked out, we obtained an image-based multi-parametric readout of how this knockout affected multiple features of a *Cryptosporidium* infection.

**Figure 1:**
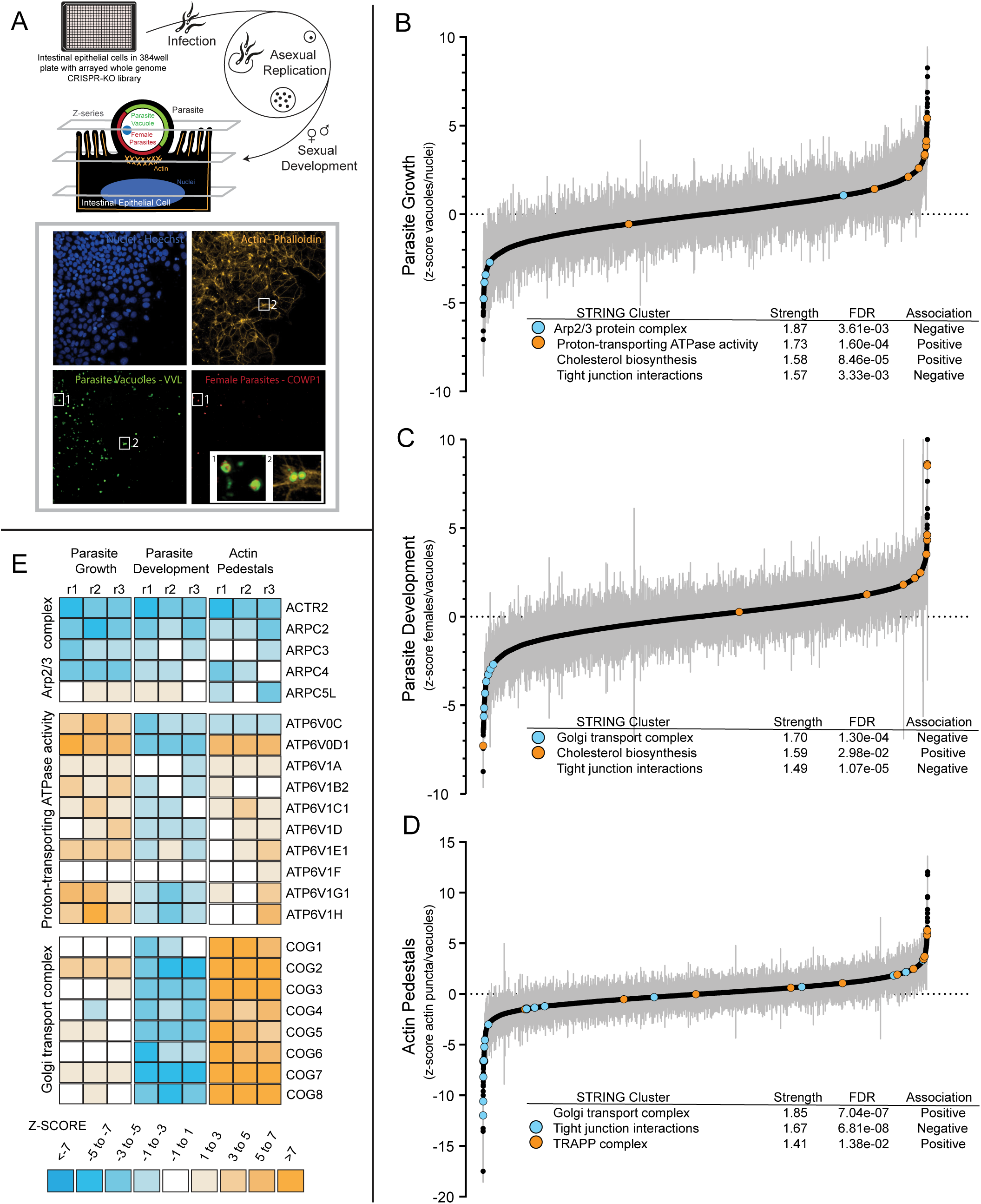
An arrayed full-genome CRISPR-Cas9 screen reveals host gene dependencies for multiple parameters of a *C. parvum* infection. **(A)** Schematic of the arrayed full-genome CRISPR-Cas9 screen. HCT-8 cells expressing Cas9 were seeded into wells containing guide crRNAs. Each well contained four guides targeted to a single gene in the human genome. Three days later, cells were infected with *C. parvum*, followed by fixation 49 hours post-infection. Cells were immunostained to detect all parasites, females, host actin, and nuclei and then imaged at three z-stacks 3 μm apart to effectively capture host nuclei, actin pedestals beneath parasite vacuoles, and apically located intracellular *Cryptosporidium*. **(B)** Rank-ordering of all 18,826 genes in the screen based on the effect of their knockout on parasite growth, **(C)** development of the female sexual stages, and **(D)** recruitment of host actin to “pedestals” beneath the parasite vacuole. Rank-ordering was based on median z-scores of three replicates per gene. **(E)** Representative examples of groups of genes in the same protein complex or pathway with significant z-scores in at least one parameter. Z-scores for each replicate in each parameter depicted as a heatmap.

To make the best use of this data in the context of a *Cryptosporidium* infection, we narrowed our primary analyses to four infection hallmarks: i) parasite growth (a ratio of the number of parasite vacuoles/number of host nuclei), ii) parasite development (a ratio of female parasites to total parasites, iii) parasite vacuoles with an associated actin pedestal, and iv) host cell viability (number of host nuclei). When we rank-ordered host genes by their median robust z-scores (measure of deviations from the median), we found a wide range of genes for which Cas9-assisted knockout either reduced or promoted parasite growth, parasite development, or actin recruitment to parasite vacuoles (Figs 1B-D). At steady state, the percentage of parasite vacuoles with detected actin pedestals slightly increases with the multiplicity of parasite infection (Fig S1G), while the percentage of female parasites in the population at 49 hours post-infection (hpi) remains constant even as the amount of infection increases (Fig S1G). This is in line with previous observations that *Cryptosporidium* sexual commitment is likely independent of parasite density^2^.

Next, we performed functional clustering analysis (via STRING) of genes that produced a significant z-score in at least one parameter^16^. Similar to transcriptomic results from genetic perturbation screens such as Perturb-seq^17^, we noticed that groups of genes whose proteins exist in complexes, or function within the same biological pathways, often produced the same phenotype across multiple infection parameters (Fig 1E). For example, loss of six out of the seven members of the actin-nucleating Arp2/3 complex, required for creation of branched F-actin filaments, reduced parasite numbers, development to female stages, and the formation of actin pedestals. Similarly, the loss of all 8 genes that function in the conserved oligomeric Golgi (COG) complex, that functions in intracellular transport, led to a uniform reduction in parasite development and an increase in actin pedestal formation during infection. Looking beyond groups of genes to the level of infection parameters, we found interesting correlations emerging between some of our selected image-based phenotypes (Fig S1K). For example, there was a significant positive correlation (r^2^ = 0.66) between host cell viability and parasite sexual development, yet no correlation between host cell viability and parasite growth (r^2^ = -0.12). This suggests that progression to sexual stages is more reliant on the health of the host cell than asexual replication.

### *Cryptosporidium* growth and development hinges at a metabolic tipping point in the host cholesterol pathway

*Cryptosporidium* lacks the ability to synthesise cholesterol and is thought to be reliant on the host to provide this essential metabolite. Therefore, one would predict that modulating this pathway would affect parasite growth and development. However, we were surprised to find that genes from the host cholesterol biosynthesis pathway were represented on either side of the *Cryptosporidium* growth and development spectra (Fig 1B, Fig 1C). While the loss of some genes in this pathway inhibited parasite growth and sexual development, the loss of others enhanced these parameters. When we plotted the z-scores for each gene in the pathway going from acetyl-CoA to cholesterol, it revealed a mid-way point at which gene loss flipped from being inhibitory to parasite growth and development, to promoting it (Fig 2A, Fig 2B). This tipping point occurred between farnesyl-diphosphate farnesyltransferase 1 (*FDFT1*), whose loss reduced *Cryptosporidium* growth and development of females, and squalene epoxidase (*SQLE*), the loss of which enhanced growth and the percentage of females.

**Figure 2:**
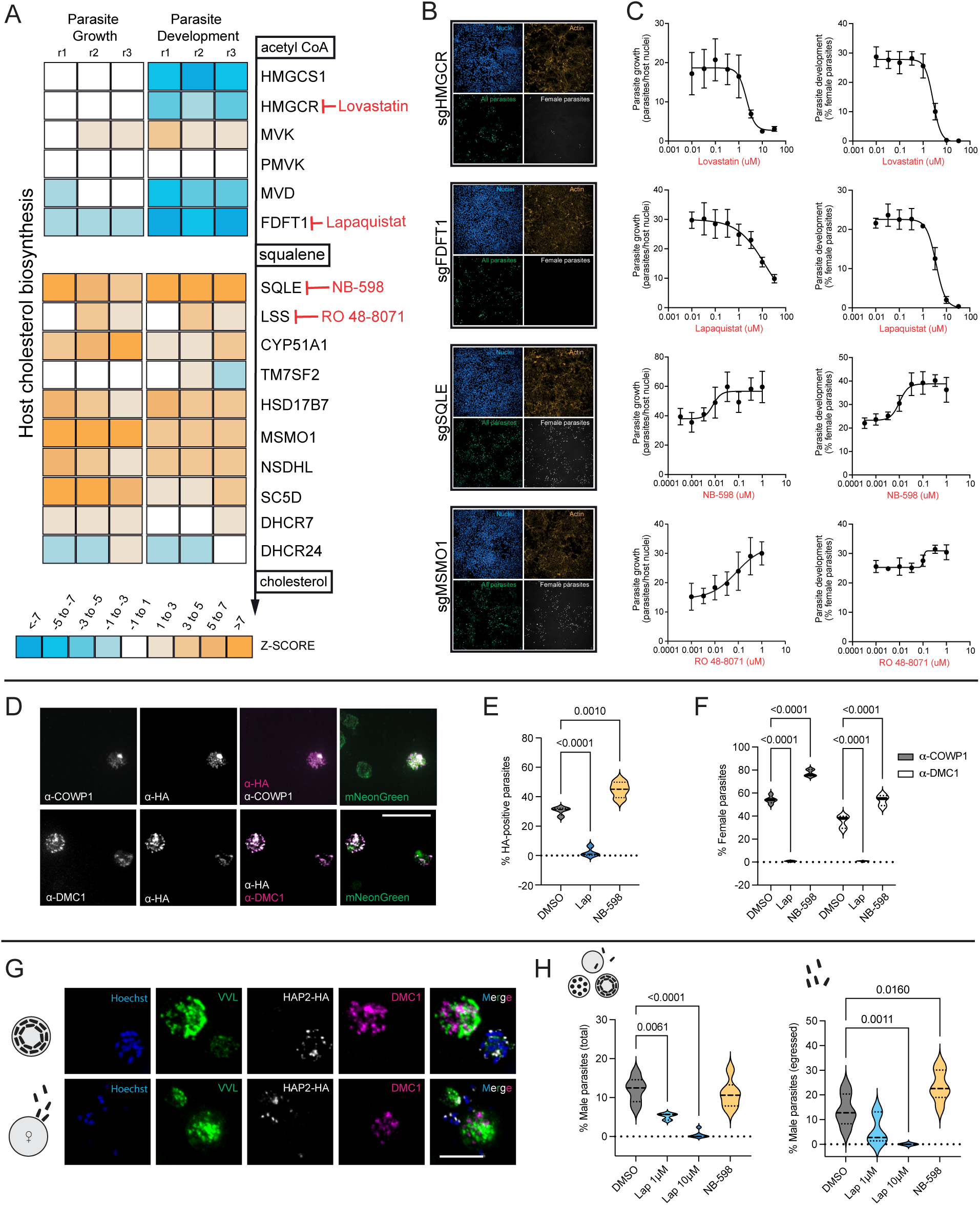
*Cryptosporidium* replication and sexual stage development hinges on a tipping point in the host cholesterol pathway. **(A)** Heatmap of z-scores displaying the effect of knocking out genes in the host cholesterol biosynthesis pathway on parasite growth and sexual development, showing a shift at the midway point between *FDFT1* and *SQLE*. Z-scores for all three replicates per gene are shown. **(B)** Representative images from the microscopy-based CRISPR screen for indicated genes in the cholesterol biosynthesis pathway showing a loss (for *HMGCR* and *FDFT1*) and increase (for *SQLE* and *MSMO1*) in total parasite numbers and females. **(C)** Dose-response curves using chemical inhibitors of indicated genes in (A), showing agreement with gene KO data in their effects on parasite replication (left) and female development (right). Parasite growth calculated as the number of parasites per host nuclei expressed as a percentage, and female development represented as the percentage of females in the total parasite population determines by anti-COWP1 staining. Data shown are the mean and SD of 6 culture wells per condition and represent one of three experiments each for lovastatin, lapaquistat, and NB-598 and two experiments for RO 48-8071. **(D)** Immunofluorescence images of HCT-8 cells infected with transgenic *Cryptosporidium* parasites expressing mNeonGreen and COWP1 endogenously tagged with HA (expressed only in females). Anti-HA antibody staining agrees with the anti-COWP1 antibody used in the screen (top row), and anti-DMC1, another female-specific antibody (bottom row). Scale bar = 5 μm. **(E)** Treatment of HCT-8 cells infected with COWP1-HA-expressing parasites with lapaquistat or NB-598 respectively reduces and increases the percentage of HA-expressing parasites 49 hpi. Data shown for 4 culture wells per condition. **(F)** Female parasite development in HCT-8 cells with indicated drug treatments identified using either anti-COWP1 or anti-DMC1 antibody staining. The percentage of parasites that were DMC1-positive is slightly lower than COWP1-positive parasites. This is likely because DMC1 is transcribed later than COWP1 during female development, hence only females that are later along in their development would be DMC1 positive^44^. Data shown for 6 culture wells per condition. **(G)** Immunofluorescence images of HCT-8 cells infected with transgenic *Cryptosporidium* parasites expressing HAP2 endogenously tagged with HA (expressed only in males). Anti-HA staining identifies bullet-shaped male microgametes at the 16-nuclei stage (top row) and egressing males finding DMC1-positive females (bottom row). Scale bar = 2 μm. **(H)** The percentage of total male parasites (left) and egressed male parasites (right) in HCT-8 cells with indicated drug treatments. Data shown for at least 3 culture wells per condition from two independent experiments. Violin plots depict the median value at the heavy dashed line and the lower and upper quartiles at the lighter dashed lines. P-values calculated using one-way ANOVA for (E), (H) and two-way ANOVA for (F) using Dunnett’s multiple comparisons test.

A previous drug repurposing screen found statins, which act on HMG-CoA reductase (HMGCR) to be a reliable host target for reducing *Cryptosporidium* growth *in vitro*^18^, so it was encouraging to see *HMGCR* coming up as a hit in our screen. We repeated this experiment using the HMG-CoA reductase inhibitor lovastatin, and we further validated our screen results by using commercially available drugs targeting enzymes on either side of the *Cryptosporidium* growth divide (Fig 2C). This revealed that lapaquistat, which targets FDFT1, reduced parasite growth and development to females in a dose-dependent manner. NB-598, which inhibits host SQLE, recapitulated the *SQLE*-knockout phenotype of enhancing growth and development, as did Ro 48-8071, which targets lanosterol synthase (LSS). Next, we were interested in which stage of infection was affected by our inhibitors. After invasion of host cells by individual parasites, they undergo many rounds of nuclear doubling within their apical vacuoles to progress from single nuclei (1n) to 2n-, 4n-, and finally 8n-parasites. At this stage, these 8 infectious forms develop and leave their vacuoles to invade neighbouring cells, repeating the asexual replication process a few more times before progression to male and female sexual stages at 36 hours post-infection. Inhibitors could be acting at any step along this route. We found that pre-treatment of host cells with lapaquistat or NB-598 did not affect parasite invasion into host cells. (Fig S2A). While lapaquistat did not affect the overall proportions of different parasite life stages at 24 hpi, it significantly reduced the numbers of 1n and 8n parasites seen per field of view in a dose-dependent manner (Fig S2B, Fig S2C). At 50 hpi, as expected, no female parasites were present at higher concentrations of lapaquistat, while the percentage of early-stage 1n, 2n, and 4n parasites increased. Therefore, while the detrimental effect of lapaquistat on *Cryptosporidium* sexual stage development is most pronounced, the drug appears to also stall asexual parasite development prior to sexual stage commitment. As NB-598 targets SQLE, the enzyme immediately following the target of lapaquistat, we wondered whether we could dampen the parasite growth-enhancing action of NB-598 with lapaquistat. Indeed, we found that increasing concentrations of lapaquistat correspondingly reduced the positive effects of NB-598 on parasite growth (Fig S2D).

To further verify the influence of the host cholesterol biosynthesis pathway on parasite development, we created a transgenic parasite line where the endogenous female specific gene *COWP1* (*Cryptosporidium* Oocyst Wall Protein 1) is C-terminally fused to an HA tag, followed by constitutive expression of mNeonGreen (Fig S2E, Fig S2F). COWP1 antibodies co-localised with antibodies targeting the HA tag (Fig 2D, top row), and lapaquistat treatment nearly eliminated the presence of HA-positive parasites (females), while NB-598 enhanced them (Fig 2E). Furthermore, another female-specific antibody for *Cryptosporidium* raised against the meiosis-specific marker DMC1^19^ was able to specifically label COWP1-HA positive parasites (Fig 2D, bottom row). DMC1 staining of infected cell monolayers revealed a similar boost from NB-598 treatment, whereas female parasites were nearly undetectable following treatment with lapaquistat (Fig 2F).

Next, we wanted to know if male parasites were similarly affected by these inhibitors. We created a transgenic parasite line where the male-specific gene *HAP2*^1^ was fused to an HA tag (Fig 2G, Fig S2G) and treated this line with either lapaquistat or NB-598. As with females, lapaquistat reduced overall male parasite development in a dose-dependent manner (Fig 2H, left). At higher concentrations of lapaquistat no females or males were observed. While NB-598 did not increase the total percentage of males in the population, when we stratified males by development stage, we found that NB-598 enhanced egress of male parasites, which is the final stage of male development (Fig 2H, right). In summary, we found that the inhibition of two consecutive enzymes in the host cholesterol biosynthesis pathway can either reduce or increase parasite growth and their overall development to male and female sexual stages *in vitro*.

### Squalene accumulation in the host epithelial cell reduces ROS and enhances *Cryptosporidium* growth

To rule out any off-target effects of NB-598 treatment, we created an *SQLE*-knockout (*SQLE*-KO) HCT-8 cell line by Cas9-directed disruption of its genetic locus (Fig S3A, Fig S3B). Loss of SQLE expression did not affect host cell viability (Fig S3C), and *Cryptosporidium* growth and development was indeed significantly enhanced in *SQLE*-KO cells compared to WT controls (Fig 3A). Differential expression analysis of transcriptomes from WT and *SQLE*-KO cells infected by *Cryptosporidium* revealed an upregulation of host cholesterol biosynthesis genes in *SQLE*-KO cells compared to WT (Fig 3B). Expression of cholesterol biosynthesis genes is tightly regulated by levels of cholesterol in the cell, sensed by sterol-responsive transcription factors such as the SREBPs which enhance transcription of cholesterol synthesis genes when cholesterol levels are low^20,21^. SQLE is a key enzyme in the cholesterol synthesis pathway; its loss would reduce cholesterol production, thereby activating transcription of cholesterol synthesis genes, which is what we found. *Cryptosporidium* parasites in *SQLE*-KO cells showed an upregulation of genes associated with late-stage female parasites compared to WT cells (Fig 3C), while the most upregulated parasite genes in WT cells by comparison were associated with late meronts, the final asexual life stage (Fig S3D). To pinpoint the consequences of changes to the host cholesterol biosynthesis pathway between WT and *SQLE*-KO cells, we next used a targeted metabolomics approach. Lovastatin treatment reduced host cholesterol levels as expected, as did both lapaquistat and NB-598 (Fig 3D, left), since they both target enzymes upstream of cholesterol production. Squalene is an intermediary metabolite in cholesterol biosynthesis, thus usually present at low levels. Treatment with either lovastatin or lapaquistat appeared to reduce squalene levels even further, as they both target enzymes upstream of squalene synthesis, while conversely, NB-598 treatment produced at least a 100-fold increase in squalene levels in host cells (Fig 3D, right). *SQLE*-KO cells had similarly elevated levels of squalene.

**Figure 3:**
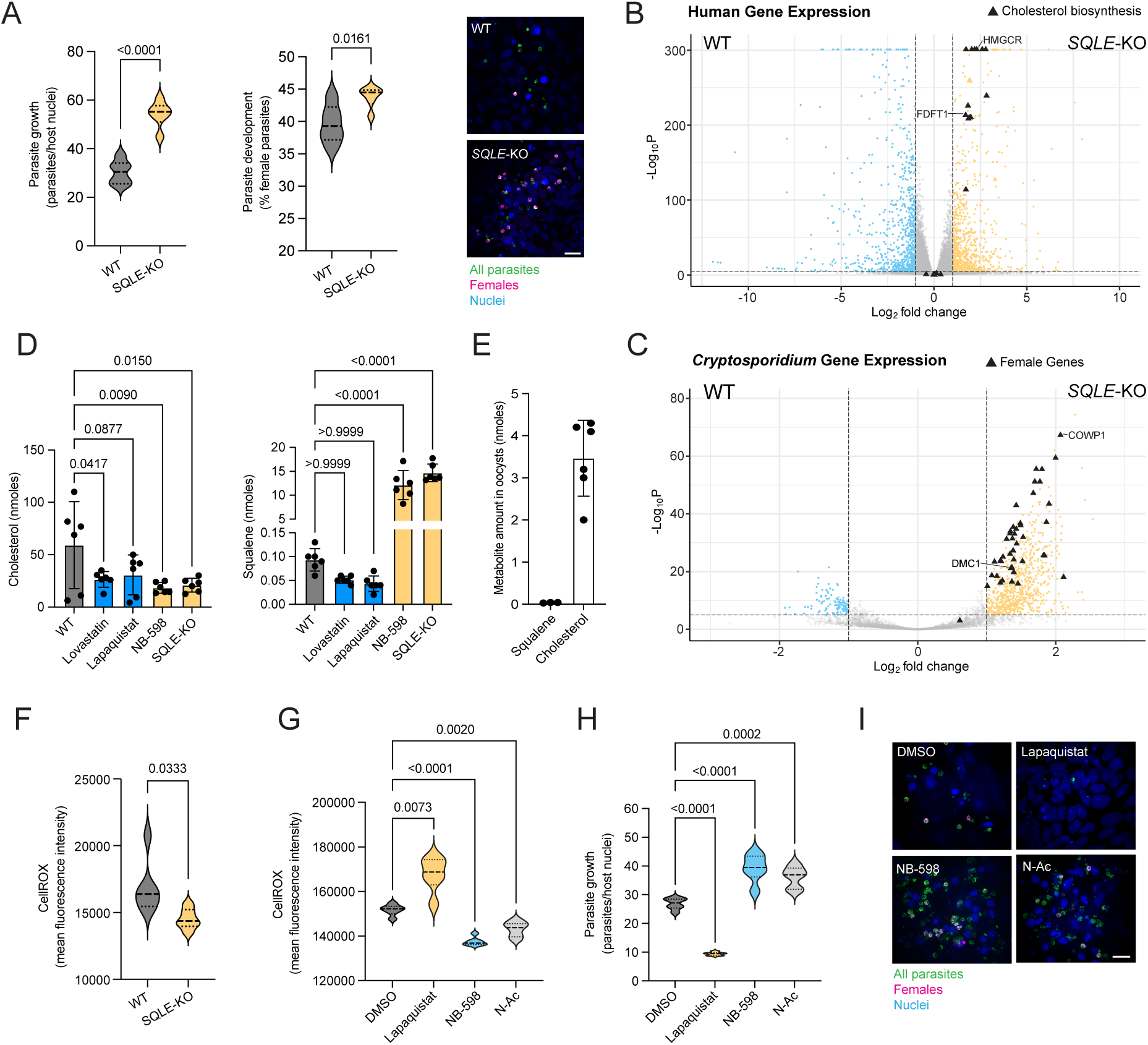
Squalene accumulation in epithelial cells reduces ROS and enhances *Cryptosporidium* growth. **(A)** Parasite replication (left), female development (middle) and representative immunofluorescence images (left) of *Cryptosporidium* infection and presence of female parasites in parental or *SQLE*-KO HCT-8 cells. Data shown for 6 culture wells per condition, representative of four independent experiments. **(B)** Differentially expressed human genes and *Cryptosporidium* genes **(C)** in infected parental vs SQLE-KO HCT-8 cells at 49 hpi, over 3 biological replicates. Horizontal dashed lines depict a p value cut-off of 1e-5 and vertical dashed lines are at a log_2_(fold change) of -1 and 1. **(D)** Quantification of cholesterol (left) and squalene (right) levels in HCT-8 cells with indicated drug treatments. **(E)** Levels of detectable cholesterol and squalene in *Cryptosporidium* oocysts. **(F)** Cellular ROS levels measured using CellROX in parental and *SQLE*-KO HCT-8 cells. Data shown for 6 culture wells per condition, representative of two independent experiments. **(G)** Cellular ROS levels measured using CellROX in infected HCT-8 cells 48 h after drug treatment. Data shown for 6 culture wells, representative of two independent experiments. **(H)** 48 h parasite growth in HCT-8 cells under indicated drug treatments calculated as the number of parasites per host nuclei expressed as a percentage. Data shown for 6 culture wells per condition, representative of three independent experiments. **(I)** Representative immunofluorescence images of wild-type *C. parvum* infection and presence of female parasites in HCT-8 cells with DMSO, lapaquistat (10 μM), NB-598 (0.5 μM), or N-Ac (10 mM). Violin plots depict the median value at the heavy dashed line and the lower and upper quartiles at the lighter dashed lines. P-values calculated by t-test with Welch’s correction for (A) and (D), and one-way ANOVA for (B), (E), and (F) using Dunnett’s multiple comparisons test.

We considered that squalene could be taken up and either stored or used by the parasite, and this could be the cause of its improved growth under conditions of squalene accumulation in the host cell. We looked for squalene in oocysts, the environmentally released infectious forms of the parasite. Squalene was not detectable in oocysts, but cholesterol was. (Fig 3E, Fig S4). Furthermore, our bioinformatic screens did not find any *Cryptosporidium* homologs for known genes that act on squalene. This suggested that while host cells supporting improved *Cryptosporidium* growth had greater levels of squalene, this metabolite does not appear to be imported or used by the parasite.

Squalene is a triterpene shown to be important for survival of B-cell lymphomas by overcoming oxidative stress^22^. We asked if squalene could be playing a similar role in epithelial cells. Cellular reactive oxygen species (ROS) levels were lower in *SQLE*-KO cells compared to WT HCT-8s (Fig 3F). Importantly, lapaquistat treatment of HCT-8 cells increased ROS within HCT-8 cells, while NB-598 which causes a squalene build-up reduced ROS levels (Fig 3G). This ROS trend was negatively correlated with parasite growth as seen previously. Interestingly, NB-598 reduced intracellular ROS to similar levels as our positive control N-acetyl cysteine (N-Ac), which is commonly used as an antioxidant. Furthermore, treatment with N-Ac alone enhanced *Cryptosporidium* growth in HCT-8 cells, comparable to NB-598 treatment (Fig 3H, Fig 3I). We also tested a range of other antioxidants on their ability to boost *Cryptosporidium* infection, finding several such as AD4 (a more membrane-permeable derivative of N-Ac), tocopherol, and ascorbic acid as being able to enhance either *Cryptosporidium* growth, sexual development, or both (Fig S5A).

### Squalene regulates levels of reduced glutathione in epithelial cells

An accumulation of the intermediary metabolite squalene was able to reduce cellular ROS and promoted parasite growth (Fig 4A). The principal mechanism whereby cells regulate ROS is via the ubiquitous antioxidant metabolite glutathione (GSH). Could increased levels of GSH alone be beneficial to *Cryptosporidium*? We first tested this by supplementing our media with glutathione ethyl ester (GSH-EE), a cell-permeable formulation of GSH. Increasing amounts of GSH-EE alone correspondingly increased parasite growth and enhanced the percentage of female parasites to similar levels as NB-598 (Fig 4B). Erastin is a small molecule that inhibits the cellular import of cystine, a key component of GSH, thus lowering the levels of intracellular GSH^23^ (Fig 4A). We found that erastin reduced parasite growth and development in a dose-dependent manner (Fig 4C). Crucially, it is effective at stopping parasite growth at concentrations that do not affect host cell viability (Fig S5B). Another inhibitor of the GSH synthesis pathway is buthionine sulfoximine (BSO), which targets gamma-glutamylcysteine synthase (GSS). BSO was similarly able to inhibit *Cryptosporidium* growth and sexual development (Fig S5C).

**Figure 4:**
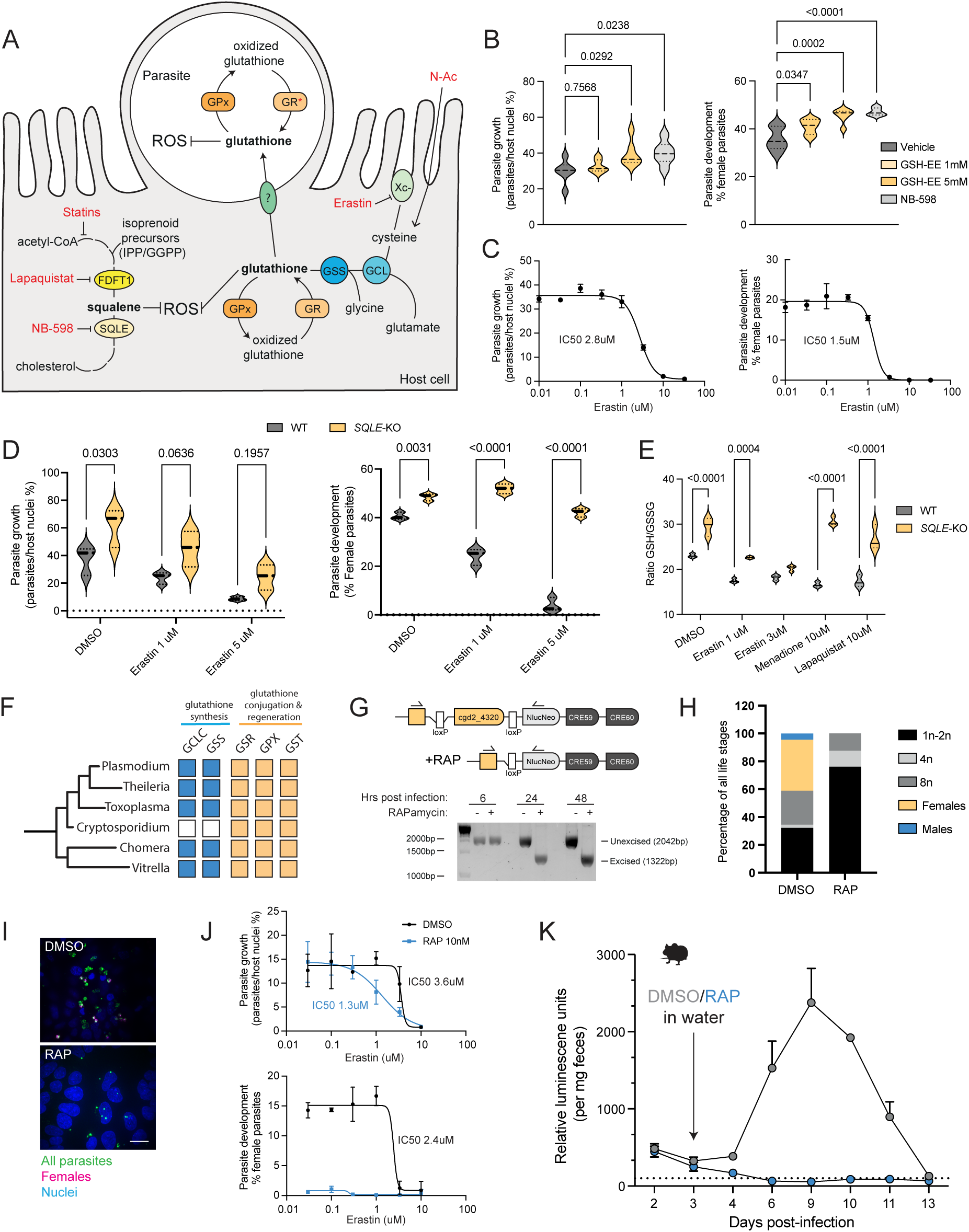
*Cryptosporidium* requires host glutathione to complete its life cycle. **(A)** Schematic of proposed model for ROS reduction by squalene and N-Ac via increased availability of reduced host glutathione for uptake by *Cryptosporidium*. Points of action of various chemical inhibitors indicated. **(B)** Parasite replication (left) and female development (right) 49 hpi with glutathione ethyl ester (GSH-EE) and NB-598 treatment. Data shown for 6 culture wells per condition, representative of two independent experiments. **(C)** Dose response curves for parasite replication (left) and female development (right) in HCT-8 cells with erastin treatment 49 hpi. Data shown are the mean and error for 3 culture wells per condition, representative of three independent experiments. **(D)** Dose response curves for parasite replication (left) and female development (right) 49 hpi in either parental or *SQLE*-KO HCT-8 cells with erastin treatment. Data shown for three wells per condition, representative of two independent experiments. **(E)** Ratio of reduced to oxidised GSH in either parental or *SQLE*-KO HCT-8 cells 4 hours after indicated drug treatments. Data shown for three wells per condition, representative of two independent experiments. **(F)** Presence of genes for GSH synthesis (mustard) or GSH recycling and conjugation (blue) in apicomplexan parasites and nearest relatives. **(G)** Schematic for the creation of transgenic *C. parvum* with loxP sites flanking the dimerization domain of its glutathione reductase gene and an inducible Cre recombinase (called *GR*-KO from here on; top). Addition of 10 nM rapamycin causes an excision of the loxP-flanked region of the parasite *GR* gene. Immunofluorescence images of HCT-8 cells infected with the inducible *GR*-KO parasites with or without rapamycin treatment (below). **(H)** Percentage of *GR*-KO parasites at indicated life stages in HCT-8 cells treated with either DMSO or 10 nM rapamycin 49 hpi. (90 parasites counted for the DMSO condition and 88 parasites counted for rapamycin condition). **(I)** Representative immunofluorescence images of inducible GR-KO parasite infection and presence of female parasites in HCT-8 cells with or without rapamycin (RAP) treatment. **(J)** Dose-response curves for replication (top) and female development (bottom) of *GR*-KO parasites in HCT-8 cells under erastin treatment with or without 10 nM rapamycin to induce loss of parasite *GR*. Data shown are the mean and error for three culture wells per condition. **(K)** Fecal nanoluciferase readings from *IFN*γ*R*^-/-^ mice infected with inducible GR-KO parasites over two weeks. This transgenic parasite line also carries a Nanoluciferase (Nluc)-expressing gene so parasite burdens can be assessed in mice by measuring fecal Nluc readings. Mice were infected by oral gavage and then either given DMSO or rapamycin in their drinking water 3 days post-infection; 2 mice per condition. Violin plots depict the median value at the heavy dashed line and the lower and upper quartiles at the lighter dashed lines. P-values calculated by one-way ANOVA using Dunnett’s multiple comparisons test (B) and two-way ANOVA using Šídák’s multiple comparisons test (D) and (E).

We next asked whether the pro-oxidative effects of erastin could be countered by the antioxidant effects of squalene. WT or *SQLE*-KO cells were infected with *Cryptosporidium* and treated with increasing amounts of erastin. While growth and sexual development in WT cells faltered early with lower levels of erastin treatment, parasites grown in *SQLE*-KO cells were much more resistant to erastin (Fig 4D). Modulators of host ROS via GSH have also been implicated in a specialised form of cell death mediated by lipid peroxidation known as ferroptosis^24^. We tested several reported inducers of ferroptosis (Fig S5D-F) and while parasite growth was inhibited, these compounds also greatly reduced host cell viability, confounding any possible interpretations of the results.

Finally, as a build-up of squalene was able to enhance parasite growth and development to similar levels as GSH-EE alone, we hypothesised that squalene may act by maintaining higher levels of reduced cellular GSH. To test this, we measured the ratio of reduced to oxidised glutathione in WT and *SQLE*-KO cells, with different inducers of ROS. *SQLE*-KO cells maintained significantly higher levels of reduced:oxidised glutathione (GSH:GSSG) in response to various ROS inducers, including lapaquistat and known cellular GSH depleters erastin and menadione (Fig 4E). Thus, increased levels of squalene in epithelial cells can buffer against GSH oxidation in the event of ROS induction, thereby maintaining a higher cellular pool of reduced GSH that is available for *Cryptosporidium*.

### *Cryptosporidium* requires host glutathione for completion of its life cycle

How might host GSH be beneficial to *Cryptosporidium*? Interestingly, unlike most other eukaryotes as well as related parasites in its apicomplexan family, *Cryptosporidium* does not possess the genes to synthesise its own GSH (Fig 4F). It does, however, possess the genes to recycle GSH from its oxidised to reduced states, as well as glutathione S-transferases (GSTs) which use GSH to post-translationally modify other proteins. Consequently, while it cannot make its own GSH, the possession of these GSH-utilising genes implies *Cryptosporidium* uses this essential metabolite and must obtain GSH from its host cell.

To test the importance of GSH for parasite fitness, we chose to impair *Cryptosporidium*’s capacity to recycle oxidised GSH by targeting its single *glutathione reductase* (*GR*) gene (cgd2_4320) for controlled disruption using an inducible gene knockout system (Fig 4G). Briefly, the presence of rapamycin brings two subunits of a Cre recombinase together, enabling excision of the loxP-flanked dimerization domain of the parasite’s *GR* gene. In the presence of rapamycin, the loxP-flanked domain is still intact 6 hpi but is completely excised 24 hpi (Fig4G). Hence, in rapamycin-treated *GR*-KO parasites, we predict loss of GSH reductase gene activity would be complete by the parasite’s third asexual replication cycle, and before its progression to sexual development. Rapamycin treatment of infected cells alone does not affect a wild-type *Cryptosporidium* infection *in vitro* (Fig S6A), however the addition of rapamycin to an infection with this transgenic strain abolished the development of male and female parasites, and skewed asexual parasites towards smaller, earlier stages of development (Fig 4H-I). While the overall numbers of parasites were not reduced, their asexual and sexual development trajectories were impacted (Fig S6B-D). We conclude that removing the parasite’s ability to recycle GSH stalls parasite growth and prevents sexual stage progression, similar to the effects of lapaquistat.

As erastin decreases host GSH levels, we tested the impact of increasing concentrations of erastin on the inducible *GR*-KO line. KO parasites were more sensitive to erastin, with their growth IC50 reducing almost 3-fold (Fig 4J). As loss of GR alone completely prevents sexual stage development, no further reductions in female parasite numbers could be assessed with erastin treatment. Finally, we tested the consequences of GR loss on a *Cryptosporidium* infection in mice. We infected two cohorts of IFNγ-deficient mice with inducible *GR*-KO parasites and administered either rapamycin or DMSO in their drinking water 3 days post-infection. *GR*-KO parasites in mice treated with rapamycin failed to establish a productive infection, with parasite burdens dropping below the detectable threshold three days post-rapamycin administration (Fig 4K). Collectively, these data show that GSH, a metabolite that the parasite cannot synthesise *de novo*, is essential for *Cryptosporidium* survival *in vivo*.

### Lapaquistat restricts *Cryptosporidium* infection in mice

In light of our evidence that inhibiting host HMGCR either genetically or chemically during an infection was sufficient to restrict parasite infection in cell culture, we were curious as to whether statins, being readily commercially available and already FDA-approved for lowering cholesterol, would also work to reduce *Cryptosporidium* growth in mice. Surprisingly, we found that treatment of *IFN*γ*R*^-/-^ mice with atorvastatin did not reduce the parasite burden in mice (Fig 5A). Mammalian cholesterol biosynthesis begins with acetyl-CoA, but there are intermediary metabolites that can feed into this pathway, such as isoprenoid precursors (Fig 4A). It has been shown that treatment with the isoprenoid precursor IPP (isopentenyl pyrophosphate) partially blocks the inhibitory effect of statins on *Cryptosporidium* growth *in vitro*^18^, and we arrived at a similar result. Inhibition of growth and parasite sexual stage development by statins was rescued by the exogenous addition of IPP or GGPP (geranylgeranyl diphosphate) (Fig 5B). In contrast, the inhibitory effect on growth and development of the parasite by lapaquistat, which inhibits FDFT1 downstream of isoprenoid entry in the cholesterol synthesis pathway, could not be rescued by the addition of isoprenoid precursors. This suggests that statins fail to limit parasite growth *in vivo* due to the presence of exogenous isoprenoids in the gut, which can bypass statin inhibition which occurs early in the cholesterol synthesis pathway.

**Figure 5:**
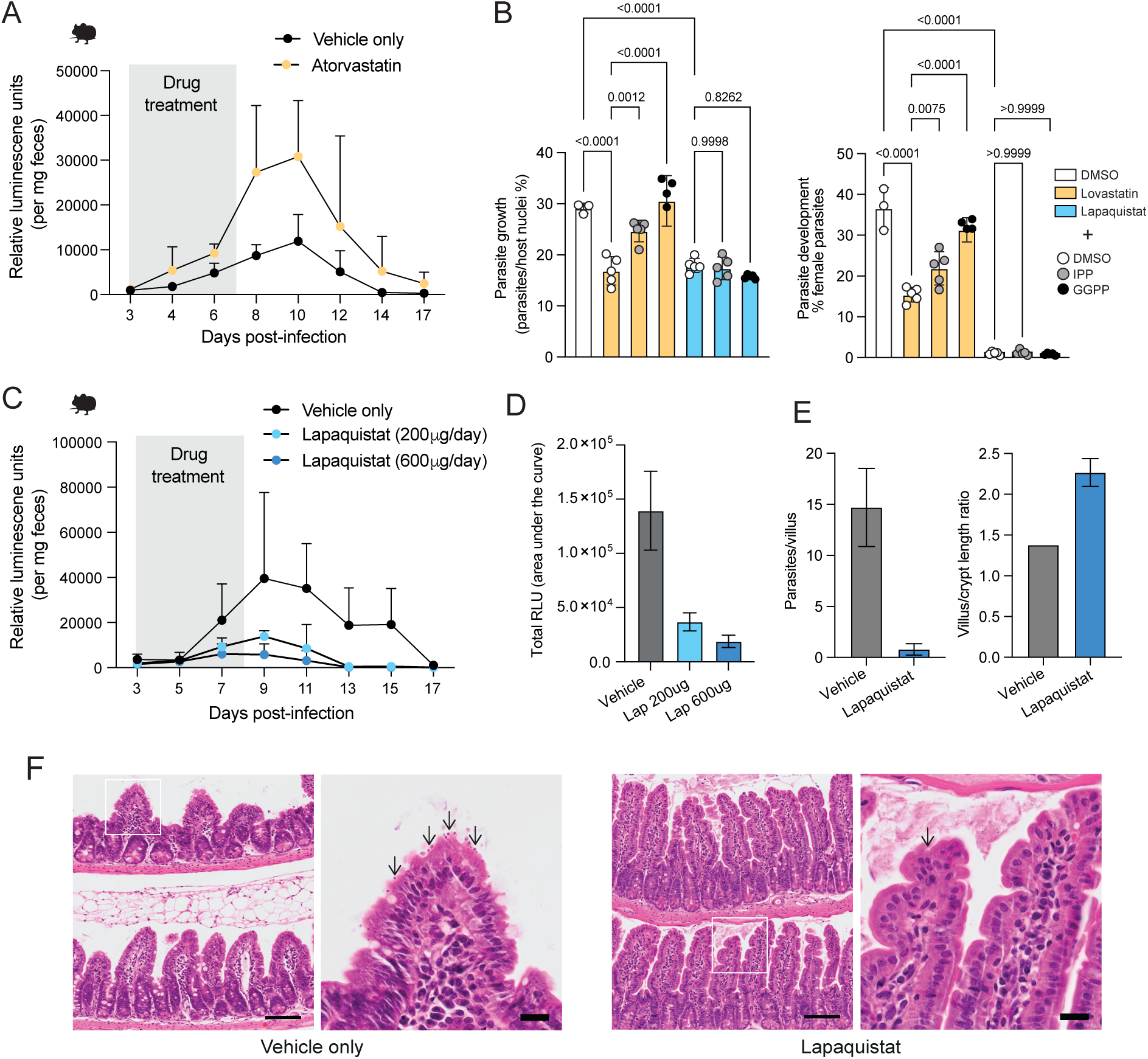
Lapaquistat restricts *Cryptosporidium* infection in mice. **(A)** Fecal Nluc readings from *IFN*γ*R*^-/-^ mice infected with a mouse-adapted *C. parvum*-mCherry line expressing Nluc. Mice were infected and given Atorvastatin (200 μg/mouse) or vehicle once daily by oral gavage on days 3-7 post-infection. 5 mice per condition; 2 mice per condition were culled on day 10 for intestinal histopathology. **(B)** Replication (left) or female development (right) of wild-type parasites infecting HCT-8 cells treated with either lovastatin or lapaquistat and additional isoprenoids as indicated. Data shown for at least 3 culture wells per condition, representative of two independent experiments. P-values calculated by two-way ANOVA using Tukey’s multiple comparisons test. **(C)** Fecal Nluc readings from *IFN*γ*R*^-/-^ mice infected with a mouse-adapted *C. parvum* line expressing Nluc. Mice were infected and given either a low (200 μg/mouse) or high (600 μg/mouse) dose of lapaquistat or vehicle only once daily by oral gavage on days 3-8 post-infection. 4 mice per condition; one mouse in the vehicle only cohort reached a humane endpoint on day 10. **(D)** Areas under the curve for Nluc levels in (C). **(E)** *C. parvum*-infected mice treated with either lapaquistat or vehicle were culled on day 10 post-infection and ileal sections were examined by H&E staining to count the number of visible parasites per villus (left) and villus-crypt heigh rations (right). 2 mice per condition. **(F)** Representative images of the ileum from (E). Scale bars are 100 μm for the main images and 20 μm for the insets.

Unlike statins, the efficacy of lapaquistat *in vitro* correlated with *in vivo* treatment. Within a murine model of acute infection, we found that treatment of infected mice, using dosage comparable to those previously trialled in human safety studies, significantly reduced parasite burden compared to vehicle-treated controls (Fig 5C). When vehicle- or lapaquistat-treated mice were culled at their infection peak, we noticed reduced numbers of parasites within the ileum of lapaquistat-treated mice (Fig 5D). Lapaquistat-treated mice also showed a decreased level of small intestinal damage that is commonly associated with *Cryptosporidium* infections, such as a decreased villus/crypt length ratio (Fig 5E, Fig 5F). Because lapaquistat already has a considerable amount of clinical trial safety data for the treatment of hypercholesterolemia, it is a highly attractive candidate for repurposing as an anti-cryptosporidial drug.

## DISCUSSION

A systematic interrogation of every protein-coding human gene and its effects on *Cryptosporidium* infection revealed unexpected insights into host and parasite cell biology, as well as a new host-directed therapeutic target for treatment of the deadly disease caused by this pathogen. We found FDFT1, the first enzyme representing commitment to cholesterol biosynthesis, to be critical for *Cryptosporidium* growth and sexual stage development. Treatment of infected mice with an inhibitor of FDFT1, lapaquistat, reduced the parasite burden and hallmarks of intestinal damage. Lapaquistat was initially developed as an alternative cholesterol-lowering drug to statins, and progressed to several phase 3 clinical trials, including one where subjects were given low or high doses of the drug (equivalent to the low or high doses given to mice in our study), both of which effectively reduced plasma LDL-cholesterol^25^. A small number of participants (two out of more than 5000) given the highest drug dose developed mildly elevated liver enzymes over a dosage period exceeding 8-12 weeks, which ultimately halted its progress to the market. This effect is believed to be on-target, as liver-specific *FDFT1* knockout mice exhibited a similar increase in liver enzymes and a build-up of isoprenoid precursors^26^. Importantly, liver enzyme levels normalised in all patients of the lapaquistat study once treatment was halted, and no long-term effects were reported. Given our finding of lapaquistat’s success in restricting *Cryptosporidium* infection in immunocompromised mice over a much shorter dosage window in this study, we are hopeful for its repurposing as an anti-cryptosporidial therapeutic. This is urgently needed since the only other FDA-approved drug for *Cryptosporidium* treatment, nitazoxanide, does not work in vulnerable populations^9,10^. Additionally, development of drug resistance is a huge barrier to fighting bacterial and parasitic diseases^27–32^ which many pathogen-targeted drug discovery studies ignore. Host-directed therapies have a much lower probability of enabling the development of resistance and are being pursued as alternative strategies against other pathogens such as *Plasmodium*^33^, *Salmonella*^34^, and SARS-CoV2^12^. Despite lovastatin and lapaquistat exhibiting synergistic potential to curb *Cryptosporidium* infection *in vitro* (Fig S7A-C), the failure of statins alone in mice and the rescue of parasite growth by isoprenoids *in vitro* cautioned us against pursuing a combined effect of the two drugs *in vivo*. We did, however, find promising *in vitro* drug efficacy with potential for additive inhibitory effects by combining lapaquistat and leading anti-cryptosporidial compounds currently in the development pipeline (Fig S7D-F).

Glutathione is the primary metabolite in the defence against oxidative damage in cells across all kingdoms of life^35–37^. Remarkably, we found that while *Cryptosporidium* carries genes to utilise GSH, in contrast to most eukaryotes, it does not have the genes for GSH synthesis. In fact, as of this writing, *Cryptosporidium* appears to be the only intracellular parasite that relies on host GSH reserves. When host GSH synthesis is blocked, via chemical inhibitors such as erastin or BSO, *Cryptosporidium* growth and development is restricted. *Cryptosporidium* has an extremely streamlined genome and carries almost 200 putative transporters to obtain various amino acids, nucleotides, sugars, and other metabolites that it cannot synthesise from its host cell^38–40^. While no definitive GSH transporters have been identified, *Cryptosporidium* gene candidates in the Major Facilitator Superfamily (MFS) of transporters with homology to a recently discovered GSH transporter in bacteria^41^ (cgd3_3100 and cgd8_3910) warrant further study. It has been shown that loss of GSH synthesis genes is detrimental to the asexual stages of related parasites *Plasmodium*^42^ and *Toxoplasma*^43^, however their importance for sexual stage development is unknown. *Cryptosporidium* expresses its *glutathione reductase* (GR) gene to recycle GSH throughout its asexual and sexual life cycle, yet two glutathione S-transferases (GSTs), enzymes that post-translationally modify proteins with GSH, appear to be expressed exclusively in female (cgd7_4780) and male (cgd8_2970) parasites^44^. The redox state, particularly GSH levels, of mouse^45,46^ and human oocytes^47^ and sperm^48^ has been shown to be important for their viability and motility respectively, with the ability to affect the rate of successful fertilisation and blastocyst formation. Our observation of greater male egress under a reducing environment aligns with this, suggesting that maintenance of proper redox balance in germ cells may be a more widespread requirement in eukaryotes.

A vast and robust dataset on *Cryptosporidium*-host dependencies and interactions generated by this whole genome KO study awaits further inquiry. Our dataset includes other parasite infection phenotypes such as the recruitment of host actin beneath the parasite, parasite vacuole size, and host viability data, revealing many other host genes that affect these aspects of *Cryptosporidium* infection when knocked out, lighting the way for follow-up studies. Interestingly, we found that host genes involved in the maintenance of epithelial cell-cell junctions such as *ZO-1*, *JAM-A* and *E-cadherin* were also important for *Cryptosporidium* growth and female development, producing a significant increase in smaller, less developed, parasites when disrupted, while also having relatively minimal effects on host cell viability. On the pathogen side, these results also allow host factors affecting sexual stage development to now be dissected away from the contexts of parasite entry and actin pedestal formation, since a negative effect on one infection phenotype did not necessarily always result in a negative effect on another. For example, knocking out components of the proton-pumping V-ATPases enhanced parasite numbers, however progression to female development was slightly reduced. V-ATPases are membrane-bound proton pumps with essential functions in basic physiology and infection, particularly in the functioning of the lysosome^49^. Lysosomes are major players in innate cellular defence against invading pathogens, representing a promising avenue of future *Cryptosporidium*-host interaction research.

Our arrayed CRISPR screen allowed us to not only discover a requirement for host GSH by *Cryptosporidium*, but also uncover the significant antioxidant properties of squalene in intestinal epithelial cells. In mice and humans, *Cryptosporidium* predominantly infects the ileum^50,51^, the most distal section of the small intestine, although a reason for this preference has been elusive. A recent transcriptomics analysis of the mouse and human small intestine found the ileum to be the site of maximal expression of cholesterol biosynthesis genes in both species^52^. Higher expression of cholesterol biosynthesis genes implies a higher flux through its intermediates, such as squalene, in the ileum. Our observations that *Cryptosporidium* survival in the intestine is influenced by these cholesterol intermediate metabolites may help to explain this ileal preference. Finally, from the host perspective, it will be intriguing to explore how the antioxidant effects of a squalene accumulation in the intestine may be used to our advantage. ROS, suppression of antioxidant pathways, or a lack of reduced GSH have been identified as instigators or components of many pathologies including inflammatory bowel disease (IBD)^53^, environmental enteropathy^54,55^ and cancer^56,57^. Modulating the redox state of the intestine by administering compounds capable of increasing squalene levels may prove to be effective against the development or progression of these pathologies. The *Cryptosporidium* parasite has a broad host range yet has evolved to infect a very specific cell type, intestinal epithelial cells, manipulating and remodelling them to support its growth and development. As we have demonstrated here, studying the interactions of this parasite with its host can be an effective and powerful tool to better understand small intestinal and epithelial cell biology.

## METHODS

### Experimental models

#### Human cell lines

HCT-8 cells were obtained from the ATCC and maintained by Cell Services at The Francis Crick Institute. HCT-8 Cas9 and HCT-8 *SQLE*-KO cell lines were generated for this study. Cells were grown at 37°C in an atmosphere with 5% CO_2_ in RPMI 1640 media supplemented with 10% fetal bovine serum (FBS), penicillin/streptomycin, and amphotericin B. During *Cryptosporidium* infections, media was changed to RPMI with 1 % FBS. Cell lines were routinely tested for the presence of mycoplasma.

#### Mouse strains

*IFN*γ^-/-^ or *IFN*γ*R*^-/-^ mice were bred and maintained by the Biological Research Facility at The Francis Crick Institute. Mice were housed in individually vented cages under a 12 h light/dark cycle with access to food and water *ad libitum*. Littermates of either sex were randomly used for experimental groups. Experiments were undertaken in accordance with UK Home Office regulations under a project license to A. S. (PP8575470).

#### Parasite strains

*Cryptosporidium parvum* oocysts (IOWA strain IIa) were purchased from Bunchgrass Farms, ID, USA. COWP1-HA-mNeonGreen, HAP2-HA, inducible GR-KO, and the mouse-adapted *C. parvum*-mCherry transgenic parasites were created and passaged using 4-8 week-old *IFN*γ^-/-^ mice.

### Method details

#### Generation of transgenic *Cryptosporidium* parasites

All transgenic parasites were created with IOWA strain IIA (Bunch Grass Farms, ID, USA) as the parental line using methods described previously^58,59^.Briefly, 2.5 x 10^7^ oocysts were induced to excyst following 1% sodium hypochlorite treatment and incubation in 0.75% sodium taurocholate in RPMI for 45 minutes at 37°C to release sporozoites. Sporozoites were transfected with a plasmid expressing Cas9 and a guide RNA targeting the desired region, along with a linear DNA template for repair of the Cas9-induced double-strand break by homology-directed repair. This linear fragment contained the tag of choice (3x HA for COWP1-HA and HAP2-HA parasite lines) or loxP-flanked recodonised genetic regions and split Cre recombinase (DiCre) genes (inducible *GR*-KO line), as well as genes for neomycin resistance and nanoluciferase expression, flanked by 50 bp regions of homology to the cut site. Transfected sporozoites were used to infect *IFN*γ^-/-^ mice by oral gavage as described previously^59^. Mice were then given paromomycin in their drinking water (16 mg/mL) to select for transgenic parasites. For details of guide RNAs and primers used for cloning the repair templates, please see Table S2.

#### Generation of inducible Cas9-expressing HCT-8 cells

A tetracycline-inducible Cas9-expressing HCT-8 cell line was generated using the Genome-Crisp^TM^ human AAVS1 Safe Harbour knock-in kit with Puromycin selection (kit no. SH016; SH100 for AAVS1 CRISPR-Cas9 clone and SH304 for AAVS1 Cas9 knock-in donor clone-TRE3G-Puro from GeneCopoeia). Briefly, sgRNA targeting the AAVS1 site on human chromosome 19 was used to initiate a Cas9-mediated double-strand break, allowing for insertion of a single copy of a tet-inducible Cas9-coding repair cassette. Positive clones were selected for by puromycin treatment, single-cell sorting, cell expansion, and confirmation by gene amplification and sequencing for correct gene cassette insertion, and immunoblotting for expression of the Cas9 protein after doxycycline treatment. Growth rates and susceptibilities to *Cryptosporidium* infection of selected positive clones were compared to the parental HCT-8 cell line to ensure there were no drastic changes in infection phenotypes (Fig S1A-S1C). A single clone (G09) was used for all screening experiments.

#### Arrayed genome-wide CRISPR-Cas9 screen

We first assessed the levels of protein knockdown that could be achieved in a 384-well plate format by using test crRNA guides against a few genes with reliable protein-directed antibodies. When we seeded Cas9-expressing HCT-8 cells (induced using 0.5 μg/mL doxycycline) along with transfection reagent Dharmafect 2 and 10 nM tracrRNA into wells containing 10 nM pooled guides against a particular gene, we found protein levels reduced by 70-80 % three days later (Fig S1D).

Cells could then be infected by *Cryptosporidium* oocysts following standard infection procedures as described previously^58^. Briefly, oocysts were treated with a 1% sodium hypochlorite solution for 5 minutes at 4°C, followed by washing once in PBS. Oocyst excystation was induced by incubating in 0.75% sodium taurocholate in RPMI for 10 minutes at 37°C. Oocysts were resuspended in 1% RPMI before addition to wells. During initial troubleshooting for setting up the screen, we analysed COWP1-expression over a 45-55-hour time window and discovered that the highest percentage of COWP1-positive parasites were present 49 hours post-infection. Moving forward, we fixed infected cells at 49 h post-infection in 4% formaldehyde. After fixing for 15-20 minutes, cells were washed in PBS, permeabilised in 0.25% Triton X-100, and blocked in 4% BSA. For convenient immunofluorescence-based analysis, we populated the four available imaging channels by staining for parasite vacuoles (using the Fluorescein-conjugated lectin *Vicia villosa* lectin or VVL), female parasites (detected with an antibody against female-specific oocyst wall protein COWP1), host-derived actin ‘pedestals’ formed beneath parasite vacuoles (detected by fluorescently-conjugated phalloidin), and host cell nuclei (using Hoechst) (Fig S1E). Specifically, we stained female parasites first using rat anti-COWP1 (1:700; created for this study) antibodies with an overnight stain at 4°C, followed by counter-staining with goat anti-rat AlexaFluor-647 (1:1000; Invitrogen), VVL-Fluorescein (1:5000; VectorLabs), and Hoechst (1:10,000; ThermoFisher). Fixed and stained plates were imaged using an Opera Phenix High Content Screening system (Revvity). Wells were imaged using a 20 x NA 0.8 air lens. Five fields of view per well were acquired. To obtain both host cell and apically-located intracellular parasite information, three z-stacks with a step size of 3 μm were taken using 405, 488, 568, and 640 nm excitation lasers paired with 450, 540, 600, and 690 nm emission filters. Image analyses were done using Harmony software v5.0 (Revvity) and detailed steps are further described in Supplementary Table S1.

We defined the ideal number of parasites needed to achieve a rate of infection that was reliably detectable by our high-content imaging platform, with room for the infection rate to significantly decrease or increase and still be detectable, while also not causing too much host cell death. To this end, we conducted titrations of each oocyst batch commercially obtained (as their viabilities often vary from batch to batch) to define these parameters. As we increased the number of oocysts, there was a corresponding decrease in host cell viability (Fig S1F). As a result, we aimed for an intermediate multiplicity of infection to allow room for a reliable detection of host factors that may increase or decrease parasite numbers without compromising host cell viability. At steady state, the percentage of female parasites in the population at 49 h post-infection remained constant while the percentage of parasite vacuoles with detected actin pedestals slightly increased as the rate of infection increased (Fig S1G). For the full genome screen, we aimed to conduct an arrayed knockout of more than 18,000 host genes – representing almost all protein-coding human genes – in triplicate. This amounted to more than 180 384-well plates for the full screen. To minimise variability and maximise confidence in each plate, we decided to include positive and negative control guide RNAs in each plate. However, as very little was known about host genes that influence *Cryptosporidium* infection, we first conducted a pilot screen with a smaller crRNA library against select genes, in the hopes of discovering appropriate controls. The pilot screen revealed some host genes that increased or decreased *Cryptosporidium* growth when knocked out (Fig S1H), from which we selected *RhoA* and *Rab8a* as our controls, as disabling these genes decreased and increased parasite growth respectively.

For the full genome screen, four crRNA guides against each gene were pooled and deposited per well, along with positive and negative control crRNAs in each 384-well plate (Fig S1I). HCT-8 cells induced to express Cas9 were then seeded into these wells, along with tracrRNA and transfection reagents, and allowed to grow for three days. Cells were then infected with *C. parvum* (to achieve a parasite/host nuclei percentage of 40% normal conditions) for 49 hours, followed by fixation, immunofluorescence labelling, and high-content imaging as described above. Triplicates of each plate were created, allowing us to calculate median z-scores for each infection parameter (outlined in Fig S1E), for each host gene. Plotting all the z-score values for the full screen for the parasite growth parameter (parasites/host nuclei), we found that our positive and negative controls cluster at either ends of the growth spectrum, our non-targeting guides clustered around non-significant z-scores (-2 to 2 exclusive), with our on-targeting guides normally distributed along this entire spectrum (S1J).

Knocking out many host genes expectedly reduced the viability of HCT-8 cells (Fig S1L). However, as we did not conduct a parallel screen without infection, we are unable to say whether the loss of host viability was only due to the importance of the gene for the viability of the cell, or whether the compounded effect of infection and host gene knockout caused cell death. As a result, we applied a z-score cut-off for cell viability before conducting downstream STRING-based analyses to avoid skewing the assessment of our other infection parameters.

#### CRISPR screen data analysis

Output from Harmony was collated into a single table and any features relating to total counts were corrected if they had fewer than five imaged fields by taking the average value per field and multiplying by five. Edge-effects were then corrected through positional normalisation per feature by taking the outer product of the row and column medians across all the plates to create an effect-matrix, scaling this by the feature mean and then multiplying the screen as a [plate, row, column] array by the scaled effect-matrix. Robust z-scores were calculated for features using the median and absolute-median-deviation of the sample wells per plate.

#### Generation of *SQLE*-KO HCT-8 cells

Five different crRNA guides (Horizon Discovery) against human *SQLE* were pooled to a final concentration of 10 nM and used to transfect Cas9-expressing HCT-8 cells along with 10 nM tracrRNA. 3 days post-transfection, cells were sorted by FACS to grow single clones, which were assessed on a build-up of neutral lipids by BODIPY-505 staining, followed by immunoblotting for expression of SQLE and sequencing of the *SQLE* gene locus to confirm Cas9-assisted partial gene deletion. Three positive clonal populations were identified and verified this way, and all downstream experiments were carried out with clone F02.

#### Immunofluorescence-based parasite growth assays

All drug-treatment assays were conducted in 96- or 384-well plates. HCT-8 cells were seeded 48 h prior to infection. Oocysts were treated as described above prior to host cell infection. For drug treatments, compounds were added 2-3 hours post-infection, except for invasion assays, in which case cells were pre-treated with the drug 16-24 h before infection. For drug synergy testing, compounds were added immediately prior to infection. Cells were fixed 6, 24, or 49 h post-infection in 4 % formaldehyde, washed in PBS, permeabilised in 0.25 % Triton X-100, and blocked in 4 % BSA. Female parasites were detected using mouse anti-DMC1 (1:1000) for 1 hour, followed by counter-staining with goat anti-mouse AlexaFluor-647 (1:1000), VVL-Fluorescein (1:5000), or *Helix pomatia* agglutinin (HPA)-AlexaFluor^TM^ 647 (1:5000) to detect all parasites, and Hoechst (1:10,000) to detect host nuclei. HPA is a lectin that, like VVL, detects α-*N*-acetylgalactosamine residues and so we use VVL and HPA interchangeably to detect parasites. Antibodies were obtained from Invitrogen unless otherwise stated. Plates were read using a BioTek Cytation 5 (Agilent Technologies) wide-field imaging reader with a 20x air objective. Four fields of view per culture well were imaged, and parasites and host nuclei were counted using its Gen5 software.

For higher-resolution imaging to stage asexual parasites and count male parasites, HCT-8 cells were seeded onto glass coverslips in 24-well plates and infected and drug-treated as previously described. Following fixation and staining as above, coverslips were mounted on slides with ProLong Gold antifade (Invitrogen) and imaged using a VisiTech iSIM for super-resolution imaging (100x or 150x oil immersion objectives, Olympus). Z-stacks of 0.2 µm step-size spanning a width of 5 µm were taken to capture the most number of parasites per field of view, and maximum intensity projections were created in ImageJ (version 2.1.0/1.53c).

To count males, infected cells were fixed at 45 hpi to ensure a mixed population of all stages were present. To quantify the percentage of all males in the population, early males (8n), late-stage males (16n) and ‘empty’ male vacuoles were counted as individual parasites, along with females and asexual stages. To quantify the percentage of egressed males in the population, egressed males were counted as individual parasites, and this number was added to the total number of parasites in the population.

#### RNA isolation and bioinformatic analyses

WT and *SQLE*-KO HCT-8 cells were infected following standard infection procedures as described previously. 1 x 10^6^ *C. parvum* oocysts were used to infect wells of a 6-well plate seeded with 6 x 10^5^ parental or *SQLE*-KO cells 48 h prior. 49 hours post-infection, RNA was recovered using the RNeasy Mini Kit (Qiagen). Total RNA was quantified by Bioanalyzer and Qubit (Agilent, Thermo). Ribosomal RNA was depleted and libraries were constructed using the Watchmaker RNA polyA (Watchmaker Genomics) according to the manufacturer’s instructions. The libraries were then pooled and sequenced on an Illumina NovaSeq X using 100bp paired-end read chemistry.

Raw sequencing reads were first assessed using FastQC (version 0.11.7) (https://www.bioinformatics.babraham.ac.uk/projects/fastqc/). Sequencing reads from each RNAseq sample were pseudo-aligned to both the predicted *C. parvum* full-length transcripts (CryptoDB-64_CparvumIOWA-ATCC_AnnotatedTranscripts) downloaded from CryptoDB^60^ (Release 64, accessed Aug 2023) and human genome transcripts (GRCh38.p14 Transcript sequences) downloaded from GENCODE^61^ (Human Release 46, accessed Sept 2024) using *kallisto*^62^ (v. 0.45.0). Briefly, the references were first prepared using *kallisto index*, after which, paired end data were pseudo-aligned to the indexed reference using *kallisto quant* with default parameters.

RNA-seq data was analysed in *R* (v.4.4.1); data exploration, filtering, and differential gene expression analyses were performed using DESeq2^63^ (v.1.46.0). Raw counts were filtered to remove low-abundance transcripts and differentially expressed transcripts were determined using a Wald test of raw counts between infected cell conditions. Transcripts were defined as being significantly differentially expressed based on fold change (log_2_(FC) > 1 or < -1) using *apeglm*^64^ and a Benjamini-Hochberg adjusted p value (Padj < 0.05). Volcano plots were used to visualise differentially expressed transcripts.

#### Cellular ROS measurements

Host cells were either infected and drug-treated for 48 hours (HCT-8 only) or uninfected (parental HCT-8 vs *SQLE*-KO). CellROX^TM^ Green or CellROX^TM^ DeepRed dyes were added to wells at a final concentration of 5 uM and incubated for 30 minutes at 37°C. Cells were then washed three times in PBS, fixed, stained with nuclear dye Hoechst and imaged using a BioTek Cytation 5 (Agilent Technologies) to obtain total fluorescence intensities and area covered to obtain mean fluorescence intensities (MFI).

#### GSH-GSSG ratio measurements

The ratio of reduced to oxidised glutathione (GSH) in Cas9-HCT8 (clone G09) and *SQLE*-KO cells (clone F02) was quantified using the GSH/GSSG-Glo^TM^ Assay kit (Promega) based on provided instructions. Briefly, cells were treated with indicated drugs in 1 % FCS RPMI. 4 hours post-treatment, media was removed, and cells were lysed in either Total or Oxidised Glutathione reagents, shaken and then treated with Luciferin Generation reagent. 30 minutes later, Luciferin Detection reagent was added to all wells and luciferase readings were taken using a BioTek Cytation 5 (Agilent Technologies) plate reader.

#### Quantification of cholesterol and squalene in HCT-8 cells and oocysts

Quantification of squalene and cholesterol in host cells or *Cryptosporidium* oocysts was performed by gas chromatography-mass spectrometry (GC-MS). For drug treatments of HCT-8 cells, 300,000 cells of either WT or *SQLE*-KO HCT-8 cells were seeded in 6-well plates. Two days later, 1% FCS RPMI media containing the drug or vehicle were added. 48 hours later, cells were processed for extraction of apolar metabolites as follows.

Briefly, media was removed and plates placed on ice and washed twice with ice-cold PBS. Cells were scraped into 250 µL chloroform:methanol:water (1:3:1, v/v) containing an internal standard (5 nmol ergosterol and 1 nmol tetracosane for sterol and squalene analysis, respectively), and incubated in a waterbath at 4°C for 1 hr with 3 x 8 minutes pulse sonication. *Cryptosporidium* oocysts were washed in ice-cold PBS before addition of the solvents and sonication as above. Samples were spun at 13,200 rpm for 10 minutes at 4°C and supernatant (metabolite extract) was transferred to new tubes and dried in a rotary vacuum concentrator. 250 µL chloroform:methanol:water (1:3:1, v/v) was added to dried samples, followed by 100 µL water (for sterols only: containing a second standard of 5 nmol lanosterol). Samples were briefly vortexed and centrifuged (13,200 rpm, 10 minutes, 4°C) to partition polar and apolar metabolites. The lower apolar phase (containing sterols and squalene) was transferred to a glass vial insert and dried once more, ready for analysis.

Data acquisition was performed using an Agilent 7890B-7000C GC-MS system in EI mode after resuspension of twice methanol-washed dried extracts in either (a) for sterols: 25 µL BSTFA + TMCS derivatisation reagent (Sigma, RT, >1 hr) or (b) for squalene, 25 µL hexane (Fisher, RT). GC-MS parameters were as follows: for sterols: carrier gas, helium; flow rate, 0.9 mL/min; column, DB-5MS+DG (30 m + 10 m × 0.25 mm, Agilent); inlet, 270°C; injection volume, 1 µl; temperature gradient, 80°C (2 min), ramp to 140°C (30°C/min), ramp to 250°C (5°C/min), ramp to 320°C (15°C/min, 6 min hold); Scan range was m/z 40-600. Data was acquired using MassHunter software (version B.07.02.1938). For squalene, parameters were as above, except: inlet, 250°C; temperature gradient, 70°C (1 min), ramp to 230°C (15°C/min, 2 min hold), ramp to 325°C (25°C/min, 3 min hold); Scan range was m/z 40-565.

Data analysis was performed using MANIC software, an in house-developed adaptation of the GAVIN package^65^. Metabolites were identified and quantified by comparison to authentic standards. All data are presented as mean ± standard deviation (SD) of 6 replicates for each condition. P-values calculated by ordinary one-way ANOVA.

#### Mouse drug treatments

4-6 week-old *IFN*γ*R*^-/-^ mice were infected with equal amounts of a mouse-adapted strain of *C. parvum* (expressing mCherry and nanoluciferase). Atorvastatin or lapaquistat were administered to mice once daily by oral gavage in a suspension of 0.5% carboxymethylcellulose as the vehicle. For mice infected with the inducible *GR*-KO parasites, rapamycin (0.05 mg/mL) was administered in their drinking water. Nanoluciferase levels in fecal samples were tracked as a proxy for parasite burden as previously described^50^ using a NanoGlo Luciferase kit (Promega).

## Acknowledgements

We would like to thank the Scientific Technology Platforms (STPs) at The Francis Crick Institute, particularly High Throughput Screening for their collaboration to develop the arrayed CRISPR-Cas9 screen, and the Biological Research Facility for breeding and maintenance of mouse lines. We thank Katarzyna Sala for experimental and administrative support at the start. We also greatly appreciate equipment and services provided by the Advanced Light Microscopy, Genomics, Metabolomics, Experimental Histopathology, and Cell Services STPs. Thank you to Dr. Chris Huston for the DMC1 antibody-producing hybridoma cell line and VEuPathDB for maintenance of critical databases. This work was enabled by The Francis Crick Institute, which receives its core funding from Cancer Research UK (CC2063), the Medical Research Council (CC2063), and the Wellcome Trust (CC2063). We are also grateful to the Aldama Foundation for their gracious support.

## Author contributions

A. S., N. B. M., O. S. and M. H. conceived of and designed the arrayed CRISPR-Cas9 screen. N. B. M, O. S., L. B. and N. B. conducted and analysed experiments. T. M. analysed the transcriptomics data and S. W. analysed the imaging data. L. C. W. contributed to creation of inducible KO parasites. V. N. and J. I. M. conducted and analysed GC-MS experiments. N. B. M. and A. S. wrote the manuscript with input from all authors.

## Supplementary Figures

**Supplementary Figure 1:**
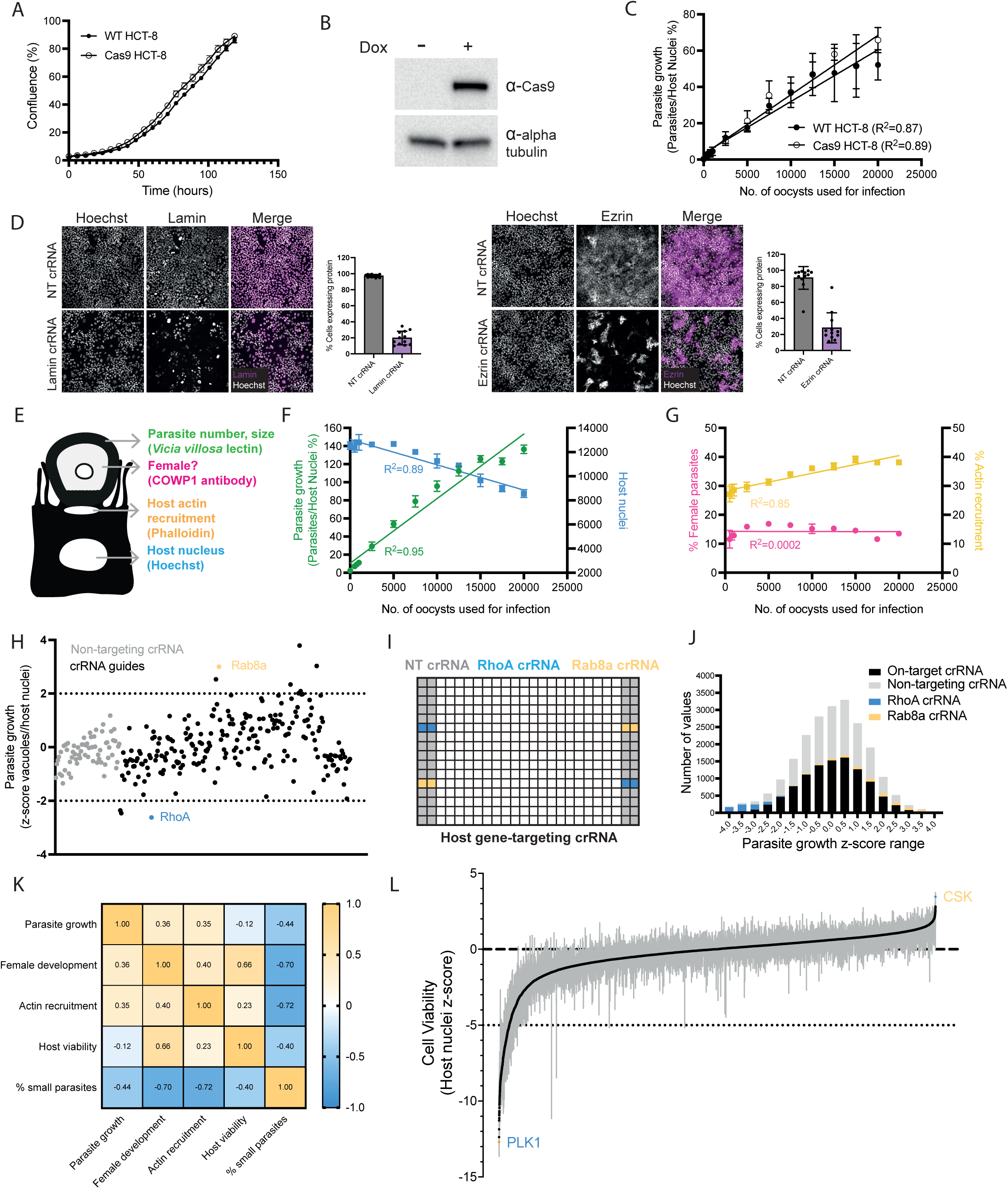
Full genome CRISPR screen setup and validation. **(A)** Growth comparison of wild-type (WT) and Cas9-HCT-8 cells measured by cell confluency using an Incucyte S3 (Sartorius) over 5 days. Mean and standard error of the mean (SEM) shown for 3 culture wells per condition. **(B)** Immunoblot of cell lysates from untreated and doxycycline-treated Cas9-HCT-8 cells to induce Cas9 expression. **(C)** Comparison of growth of *C. parvum* in WT and Cas9-HCT-8 cells. Increasing numbers of oocysts were used to infect either cell type. Parasite growth in either cell type (calculated as the number of parasites per host nuclei expressed as a percentage) was assessed 49 hpi. Mean and standard deviation (SD) of 5 wells per condition shown. **(D)** Levels of protein expression in Cas9-expressing HCT-8 cells three days post-transfection of crRNA directed against Lamin (left) or Ezrin (right), measured by immunostaining and calculating fluorescence intensities of each cell over 4 fields of view for 3 wells per condition. Scale bar = 100 μm. **(E)** Schematic showing the four parameters of a *C. parvum* infection that were targeted for immunofluorescence analysis in the CRISPR-Cas9 screen: parasite number and size, staining for female parasites, recruitment of host actin to parasite vacuoles, and the number of host nuclei. **(F)** Examination of parasite growth (green line) and numbers of host nuclei (blue line) with increasing numbers of oocysts used to infect HCT-8 cells. Note a linear relationship with initial oocyst infection numbers in both cases. Mean and SD of three culture wells per condition. Representative of three independent experiments. **(G)** Examination of the percentage of parasite vacuoles with actin pedestals (mustard line) and the percentage of female parasites in the population (magenta line) with increasing numbers of oocysts used to infect HCT-8 cells. Mean and SD of three culture wells per condition. Representative of three independent experiments. **(H)** Scatter plot of z-scores from the pilot screen for host genes affecting *C. parvum* growth, highlighting positions of *Rab8a* and *RhoA* above and below the 2/- 2 cut-off for significantly affecting growth, resulting in their selection as positive and negative controls for the full genome CRISPR screen. **(I)** Layout for each of the 183 384-well plates used in the full genome CRISPR screen. Non-targeting control guides (NT crRNA) placed in grey locations, RhoA controls in blue, Rab8a controls in mustard, and all on-target guides against host genes in white. **(J)** Distribution of parasite growth z-scores for the different control and on-target guide RNAs in the full genome CRISPR screen. **(K)** Correlation matrix of different infection parameters analysed from the screen for selected host genes with a significant z-score in at least one infection parameter. **(L)** Rank-ordering of all 18,826 genes in the screen based on their effect on host cell viability z-scores. Rank-ordering based on median z-scores of three replicates per gene. Variance for each gene shown in grey.

**Supplementary Figure 2:**
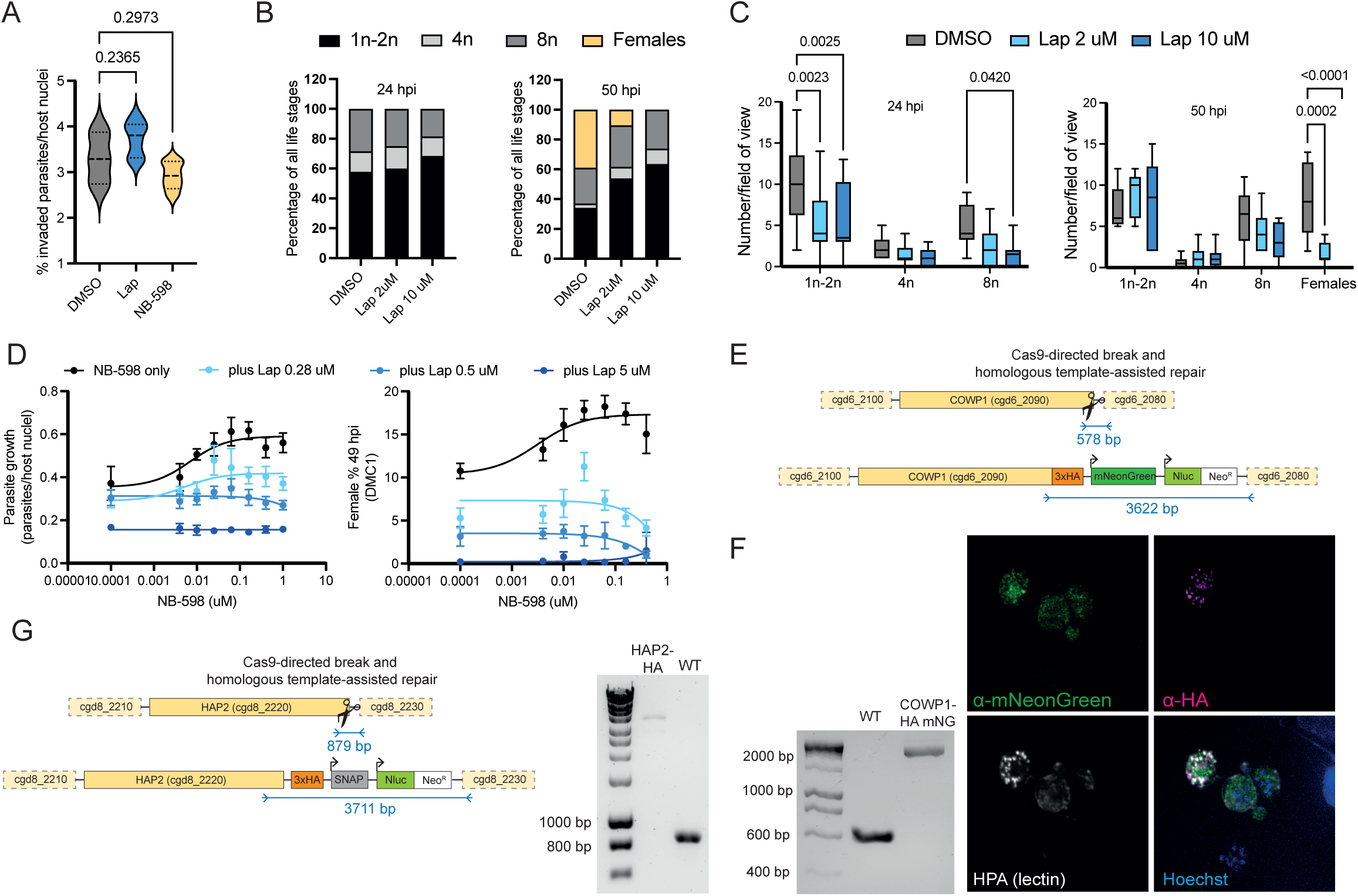
Characterising modes of action of chemical inhibitors lapaquistat and NB-598. **(A)** Lapaquistat (10 μM) and NB-598 (0.5 μM) do not affect invasion of *C. parvum* sporozoites. HCT-8 cells were treated overnight with indicated drugs and were infected with newly-excysted sporozoites the next day and fixed 4 hpi. Violin plots depict the median value at the heavy dashed line and the lower and upper quartiles at the lighter dashed lines. Data shown represent 6 culture wells per condition. P-values calculated by one-way ANOVA with Dunnett’s multiple comparisons test. **(B)** Percentage of life stages seen at 24 hpi in total (left; DMSO =173 parasites, 2 μM lapaquistat = 140 parasites, 10 μM lapaquistat = 114 parasites) and 50 hpi (right; DMSO =170 parasites, 2 μM lapaquistat = 115 parasites, 10 μM lapaquistat = 96 parasites) under drug treatment of infected HCT-8 cells. **(C)** Number of parasite life stages seen per field of view at 24 hpi and 50 hpi from (B). Mean and SD shown for at least 8 fields of view per condition. P-values calculated using two-way ANOVA with Šídák’s multiple comparisons test. **(D)** Dose-response curves for parasite growth (left) and percentage of females (right) in HCT-8 cells 49 hpi with NB-598 treatment alone (in black), or with increasing concentrations of lapaquistat (in shades of blue). **(E)** Schematic for the creation of the *C. parvum* transgenic line with endogenously tagged COWP1 by Cas9-assisted break and homology repair. **(F)** PCR-based verification of creation of the COWP1-HA-mNeonGreen parasite line (left). Genomic DNA was extracted from either wild-type parasites or purified transgenic parasites. Immunofluorescence-based verification of the COWP1-HA-mNeonGreen parasite line (right). Transgenic parasites were used to infect HCT-8s, fixed 49 hpi, and probed with an anti-mNeonGreen antibody (green), anti-HA (magenta), HPA-AlexaFluor^TM^ 647 (white), and Hoechst (blue). **(G)** Schematic for the creation of the *C. parvum* transgenic line with endogenously tagged HAP2 by Cas9-assisted break and homology repair (left). PCR-based verification of creation of the HAP2-HA parasite line (right). Genomic DNA was extracted from either wild-type parasites or purified transgenic parasites.

**Supplementary Figure 3:**
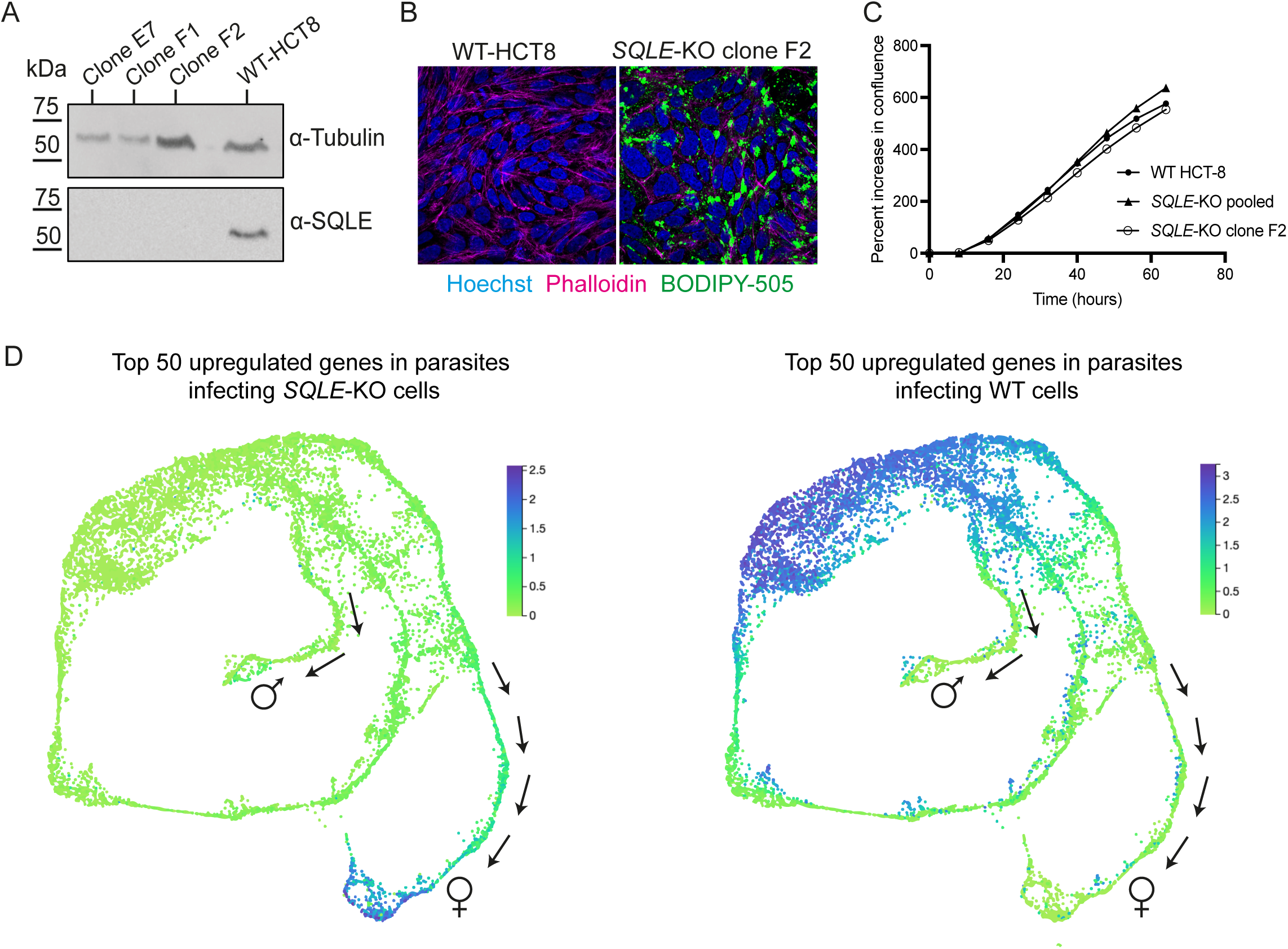
Characterisation of *SQLE*-KO HCT-8 cells. **(A)** Immunoblot for SQLE expression in wild-type (WT) HCT-8 cells and selected *SQLE*-KO clones. **(B)** *SQLE*-KO cells have a build-up of neutral lipids. Cells were seeded onto coverslips, fixed, and stained to visualise nuclei (Hoechst; blue), actin (Phalloidin-AlexaFluor^TM^ 647; magenta), and neutral lipids (BODIPY-505/515; green). **(C)** Growth comparison of WT HCT-8 cells, a mixed population of *SQLE*-KO cells, and one clonal population of *SQLE*-KO cells (clone F2) measured by the percent increase of their cell confluency using an Incucyte S3 (Sartorius) over 64 hours. Mean shown for 3 culture wells per condition. **(D)** Differentially expressed *C. parvum* genes in *SQLE*-KO and WT HCT-8 infections mapped onto the parasite life cycle. The mean expression for the 50 most significant genes (by adjusted p-value) with differentially regulated expression were mapped onto the life cycle of *C. parvum,* visually represented as a UMAP of single-cell parasite transcriptomes through CZ CELLxGENE^44,66,67^. For parasites infecting *SQLE*-KO cells, the most significantly upregulated genes cluster to late females (left). Male-specific genes were also significantly upregulated in an *SQLE*-KO infection (e.g. HAP2 had a log_2_FC of 1.6 in *SQLE*-KO vs WT cells), although to a lesser degree. Late male parasite stages are motile and not anchored to the host cell monolayer, therefore it is likely many were lost during RNA isolation. For parasites infecting WT cells, significantly upregulated genes compared to *SQLE*-KO cluster to late stage asexual meronts (right).

**Supplementary Figure 4:**
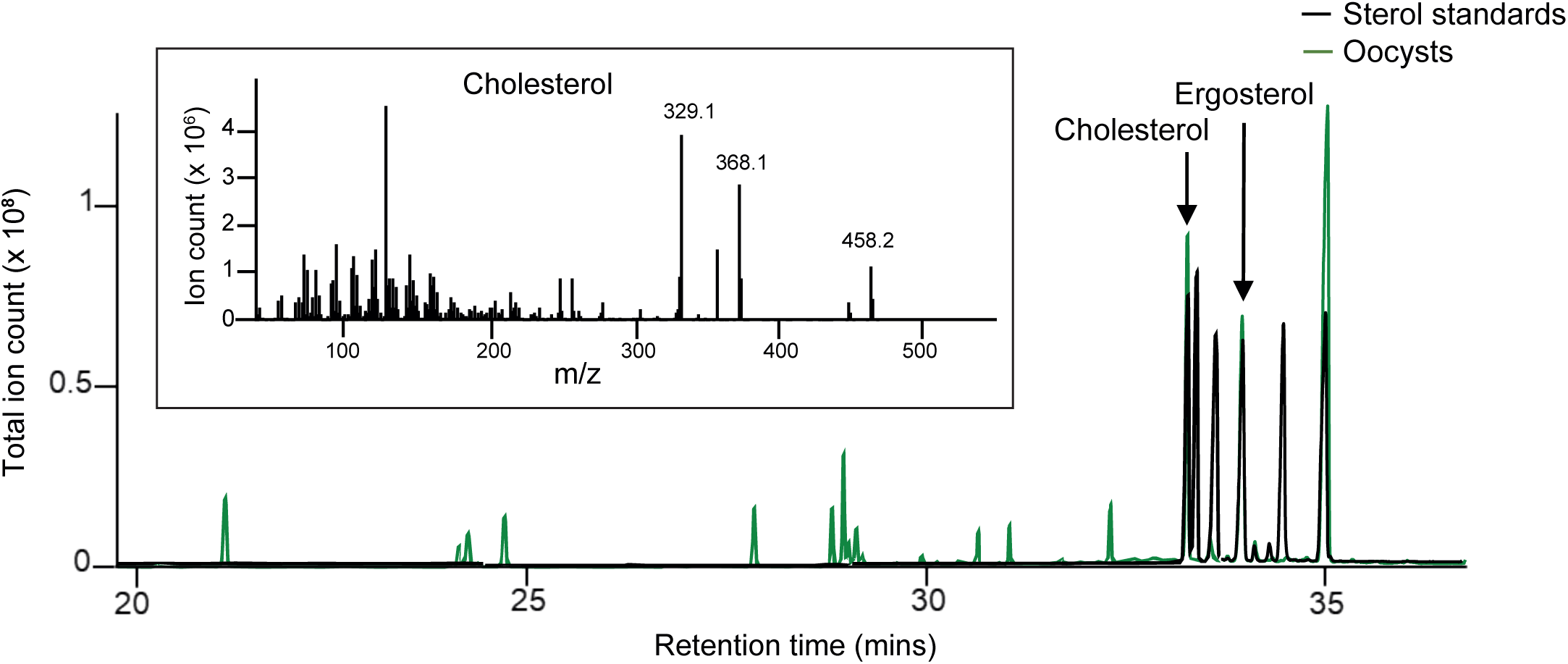
GC-MS identifies cholesterol in parasite oocysts. Gas chromatography-mass spectrometry (GC-MS) analysis of *Cryptosporidium* oocysts. Chromatograms are representative of 6 replicates. The panel shows an overlay of cholesterol analysis chromatograms of *C. parvum* oocysts (green line) and authentic sterol standards (5 nmol, black line). Cholesterol and ergosterol (internal standard) are indicated. A representative spectrum of cholesterol is shown (inset), with diagnostic ions highlighted.

**Supplementary Figure 5:**
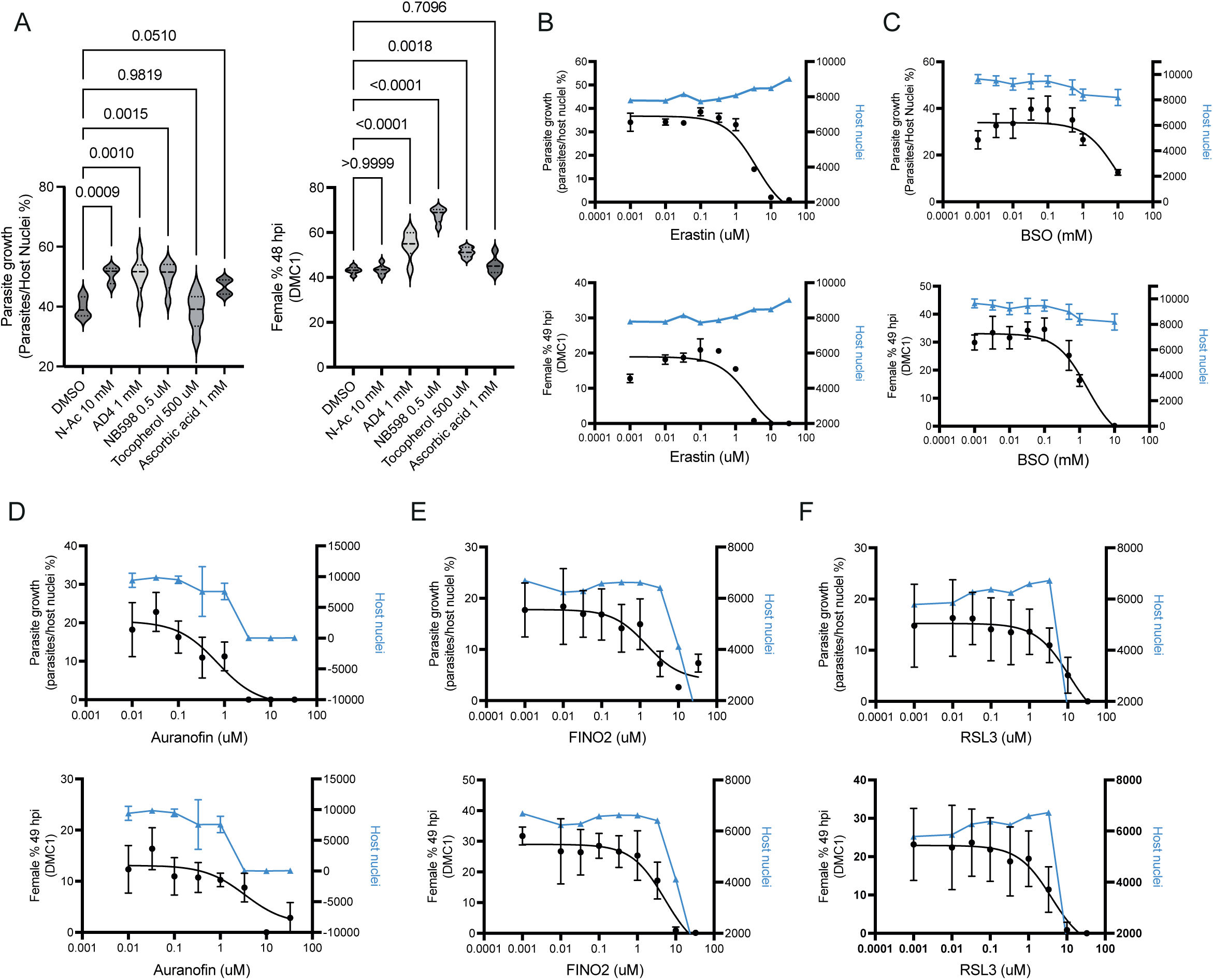
*Cryptosporidium* growth *in vitro* is affected by pro- and antioxidants. **(A)** Parasite growth (left) and female development (right) in HCT-8 cells in the presence of various antioxidants: N-Ac (10 mM), AD4 (1 mM), NB-598 (0.5 μM), tocopherol (500 μM), and ascorbic acid (1 mM). Violin plots depict the median value at the heavy dashed line and the lower and upper quartiles at the lighter dashed lines for 6 culture wells per condition. P-values calculated by one-way ANOVA with Dunnett’s multiple comparisons test. **(B)** Erastin reduces parasite growth (top) and female development (bottom) in a dose-dependent manner with a minimal effect on host cell viability (blue lines). Parasite data also shown in Fig 4C. Mean and SD for 3 culture wells shown, representative of three independent experiments. **(C)** BSO reduces parasite growth (top) and female development (bottom) in a dose-dependent manner with a minimal effect on host cell viability (blue line). Mean and SD for 3 culture wells per condition shown, representative of two independent experiments. **(D)** Auranofin reduces parasite growth (top) and female development (bottom) while also causing host cell death in a dose-dependent manner (blue lines). Mean and SD for 6 culture wells per conditions shown, representative of two independent experiments. **(E)** FINO2 and RSL3 **(F)** reduce parasite growth (top) and female development (bottom) while also causing host cell death in a dose-dependent manner (blue lines). Mean and SD for 6 culture wells per condition shown.

**Supplementary Figure 6:**
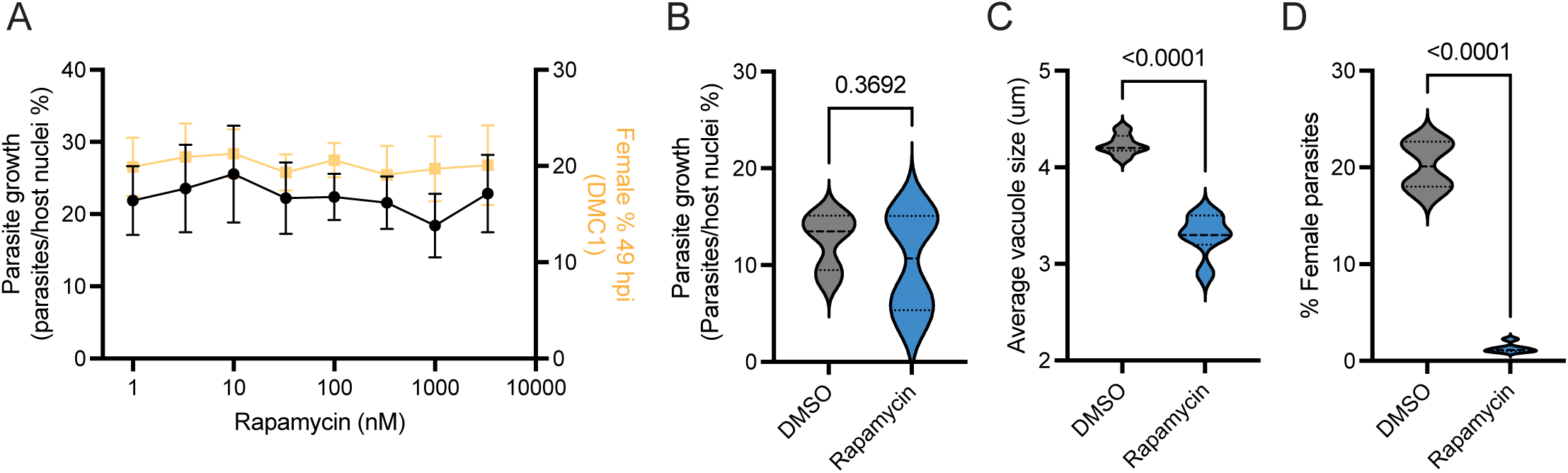
Validation of Cre recombinase-mediated loss of *Cryptosporidium* glutathione reductase (GR). **(A)** Rapamycin does not affect growth (in black) or female development (in mustard) of HCT-8 cells infected with wild-type parasites at 1 – 3000 nM. Mean and SD for 6 culture wells per condition shown. **(B)** Total *GR*-KO parasites per host nuclei, average size of all parasites **(C)**, and the percentage of female parasites **(D)** 49 hpi with or without 10 nM rapamycin treatment to induce GR loss. Violin plots depict the median value at the heavy dashed line and the lower and upper quartiles at the lighter dashed lines for 6 culture wells per condition. P-values calculated by t-tests.

**Supplementary Figure 7:**
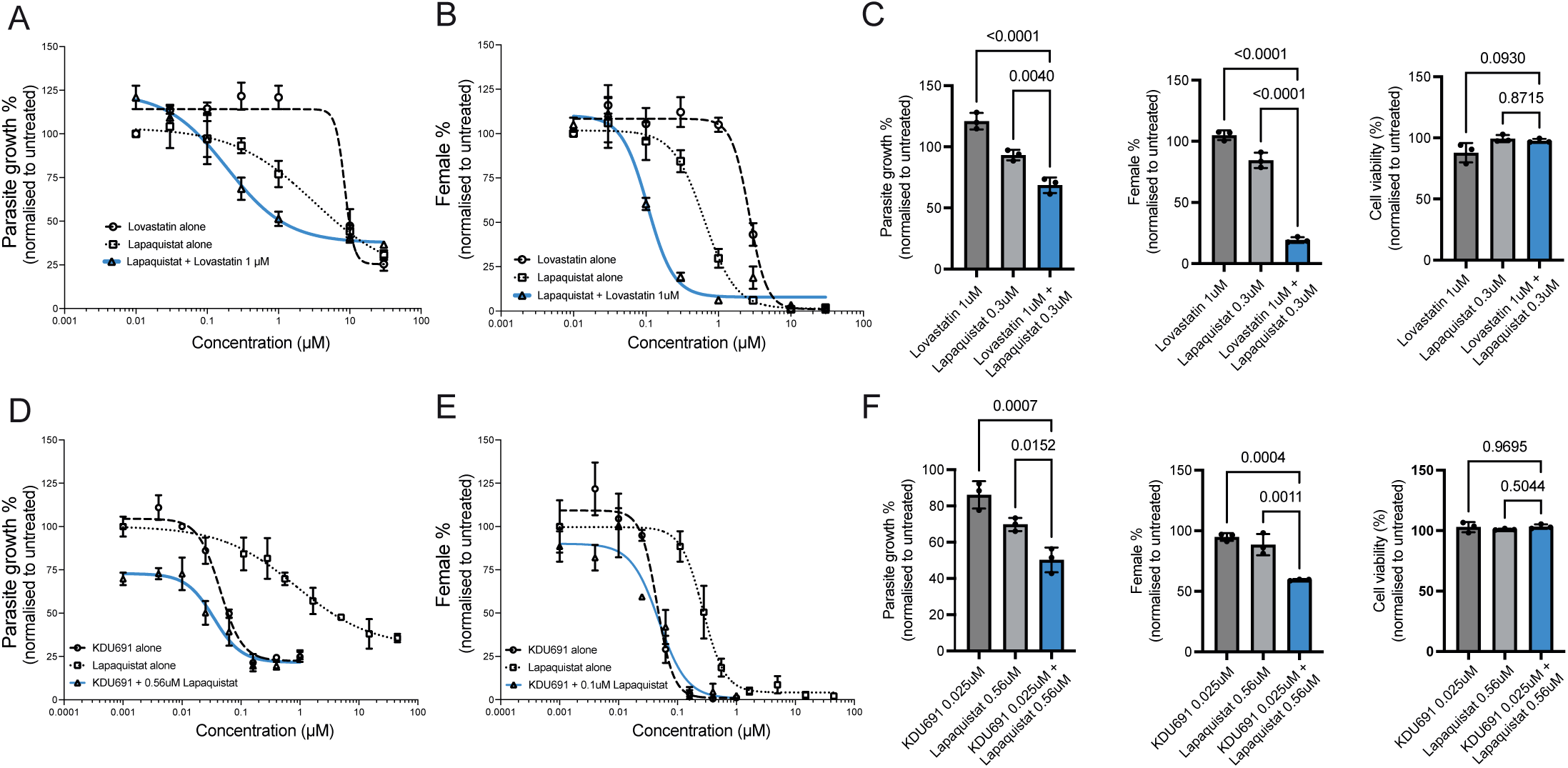
Testing lapaquistat in combination with other inhibitors for synergistic effects on *Cryptosporidium* infection. **(A)** Dose-response curves for the effect of lapaquistat or lovastatin alone, or in combination on *Cryptosporidium* growth or female development in HCT-8 cells **(B)**. **(C)** Specific combinations of lapaquistat and lovastatin at indicated drug concentrations showing synergistic effects on parasite growth and female development are shown, as well as their minimal effects on host cell viability at the same concentrations. Combination Index (CI) values of 0.2 and 0.1 for 50% inhibition of parasite growth and female development respectively, indicating synergistic effects of the two drug combinations^68^; Loewe synergy scores of 3.3 ±2.2 for parasite growth and 3.2 ±1.6 for female development^69^. **(D)** Dose-response curves for the effect of lapaquistat or KDU691, a *Cryptosporidium* PI4K inhibitor^70^ alone, or in combination on parasite growth or female development in HCT-8 cells **(E)**. **(F)** Specific combinations of lapaquistat and KDU691 at indicated drug concentrations showing additive inhibitory effects on parasite growth and female development are shown, as well as their minimal effects on host cell viability at the same concentrations. Combination Index (CI) values of 0.95 and 1.2 for 50% inhibition of parasite growth and female development, respectively, indicating additive effects of the two drug combinations; Loewe synergy scores of -3.9 ±1.4 for parasite growth and -3.4 ±3.6 for female development. P-values calculated by one-way ANOVA with Dunnett’s multiple comparisons test.

